# Molecular mechanism for regulating APOBEC3G DNA editing function by the non-catalytic domain

**DOI:** 10.1101/2024.03.11.584510

**Authors:** Hanjing Yang, Josue Pacheco, Kyumin Kim, Diako Ebrahimi, Fumiaki Ito, Xiaojiang S. Chen

## Abstract

APOBEC3G (A3G) belongs to the AID/APOBEC cytidine deaminase family and is essential for antiviral immunity. It contains two zinc-coordinated cytidine-deaminase (CD) domains. The N-terminal CD1 domain is non-catalytic but has a strong affinity for nucleic acids, whereas the C-terminal CD2 domain catalyzes C-to-U editing in single-stranded DNA. The interplay between the two domains in DNA binding and editing is not fully understood. Here, our studies on rhesus macaque A3G (rA3G) show that the DNA editing function in linear and hairpin loop DNA is greatly enhanced by AA or GA dinucleotide motifs present downstream (in the 3’-direction) but not upstream (in the 5’-direction) of the target-C editing sites. The effective distance between AA/GA and the target-C sites depends on the local DNA secondary structure. We present two co-crystal structures of rA3G bound to ssDNA containing AA and GA, revealing the contribution of the non-catalytic CD1 domain in capturing AA/GA DNA and explaining our biochemical observations. Our structural and biochemical findings elucidate the molecular mechanism underlying the cooperative function between the non-catalytic and the catalytic domains of A3G, which is critical for its antiviral role and its contribution to genome mutations in cancer.

## Introduction

Human APOBEC3G (hA3G), a member of AID/APOBEC family of zinc-containing cytidine deaminases, catalyzes the conversion of cytidine (C) to uridine (U) on DNA. This process generates DNA mutations from unrepaired uridines. hA3G is a well-known host restriction factor that plays a crucial role in restricting human immunodeficiency virus type 1 (HIV-1)^1^. In the absence of HIV viral infectivity factor (Vif), hA3G can catalyze excessive C to U editing on the HIV-1 negative cDNA strand, leading to hypermutation in HIV-1 genome^2–10^. hA3G can also impair HIV-1 replication through deaminase-independent mechanisms^11–17^.

Deaminases can also induce mutations in the host genome in the context of pathological misregulation in tumorigenesis^18–24^. Analysis of human cancers revealed hA3G’s contribution to mutational signatures in multiple cancer types^25,26^. In a murine bladder cancer model, transgenic expression of hA3G promotes mutagenesis and genomic instability^25^. Additionally, hA3G performs C to U editing on certain types of human and viral RNA^27–30^.

A3G is composed of two zinc-containing cytidine deaminase (CD) domains in tandem: the N-terminal CD1 domain or NTD (referred to as CD1 hereafter) and the C-terminal CD2 domain or CTD (referred to as CD2 hereafter). Multiple CD1-CD2 domain orientations in full-length A3G have been observed in protein crystal structures and cryo-electron microscopy (cryo-EM) structures^31–36^. Despite their similar tertiary structures, individual CD1 and CD2 domains have evolved to carry out distinct functions^37,38^. The CD1 domain is non-catalytic but binds strongly to nucleic acids^39,40^. Recent studies show that RNA purine dinucleotide sequence motifs rArA and rGrA are preferred RNA binders for the primate rhesus macaque A3G^33^. Cryo-EM studies have revealed that the rArA- or rGrA-RNA bound by A3G is a critical part recognized by HIV Vif-E3 ligase for A3G ubiquitination and proteasome degradation^34–36^. These studies provide evidence that CD1, with assistance of CD2, engages in direct RNA binding. On the other hand, the CD2 domain carries out DNA target-C editing, although it has weak affinity to DNA^4,5,41–45^. CD2 favors 3′ target-C to U editing in the motifs C<u>C</u> or CC<u>C</u> (the target-C is underlined) on single-stranded DNA (ssDNA). X-ray protein crystallography studies of the catalytic CD2 domain bound to the DNA substrate or DNA oligonucleotide inhibitor have revealed the molecular details of the editing motif CC<u>C</u> selection and deamination^44,46^.

Efficient A3G editing requires cooperativity between its two domains. The catalytic CD2 domain alone displays up to three orders of magnitude lower editing efficiency than the full-length protein^47,48^. Furthermore, full-length A3G processively edits target-C in two CC<u>C</u> motifs located on a ssDNA substrate during one binding event and preferentially edits target-C in the CC<u>C</u> motif near the 5’ end of ssDNA substrates^48–51^. These two editing properties are impaired in the absence of the non-catalytic CD1 domain^48^. Data from experiments with optical tweezers show that A3G binds in multiple steps and conformations to search and deaminate single-stranded DNA^52^. Despite these advances, the precise molecular mechanism used by the two domains to coordinate DNA binding and editing has remained elusive.

In this study, we find that purine dinucleotide AA or GA motif downstream of the target-C editing sites in the 3’-direction facilitates rhesus macaque A3G (rA3G) DNA editing function in linear and hairpin loop DNA. The effective distance between AA/GA motifs and the target-C sites depends on the local DNA secondary structure. We present two co-crystal structures of rA3G in complex with ssDNA containing AA or GA motif, providing a mechanistic understanding for AA/GA recognition through the non-catalytic CD1 domain, and its impact on the target-C selection and editing efficiency. These structures also explain RNA inhibition of DNA editing. Our findings reveal molecular insights into the cooperativity of the two domains for A3G’s efficient DNA editing, crucial for its antiviral function on foreign pathogens and mutagenic function on genomic DNA.

## Results

### Purine dinucleotide motifs facilitate DNA editing of rA3G

Previously, we have shown that rA3G has a strong binding affinity to rArA dinucleotide containing RNA with K_D_ between 10 to 17 nM, followed by rGrA dinucleotide with K_D_ of ∼47 nM, and other combinations of dinucleotide containing RNA with K_D_ of ∼124 nM or much worse^33^. It turns out that rA3G also binds AA-containing DNA (5’-FAM TTTTAATTTT) with K_D_ of ∼318 nM, and GA-containing DNA (5’-FAM TTTTGATTTT) with K_D_ of ∼473 nM (**Supplementary Fig. 1**). With this information, we hypothesized that the presence of AA or GA in DNA may facilitate substrate capturing by A3G and the presentation of nearby target-Cs to the active center on the catalytic A3G-CD2 domain.

To study whether and how the AA motifs on DNA can facilitate A3G deaminase activity, we compared editing efficiency between control DNA substrates that carry one editing motif CC<u>C</u> (the target-C is underlined) and AA-DNA substrates that carry both CC<u>C</u> and AA motifs. For simplicity, the control DNA contains only mixed pyrimidine bases, or a combination of mixed pyrimidine and guanine bases, but no adenine base. When designing single-stranded linear DNA substrates (**Fig. 1a** inset), two variables are considered: (1) substrate length and (2) distance of the editing motif CC<u>C</u> from 3’-end^48–50^. It has been reported that when distance of the editing motif CC<u>C</u> from 3’-end is less than 30 nt, the editing motif CC<u>C</u> falls into a weakly deaminated ‘dead’ zone at the 3’-end linear DNA with a specific activity of human A3G less than 1 pmol/μg/min^48,50^. We wished to study whether AA motifs could facilitate A3G editing efficiency under such situation.

**Figure 1.**
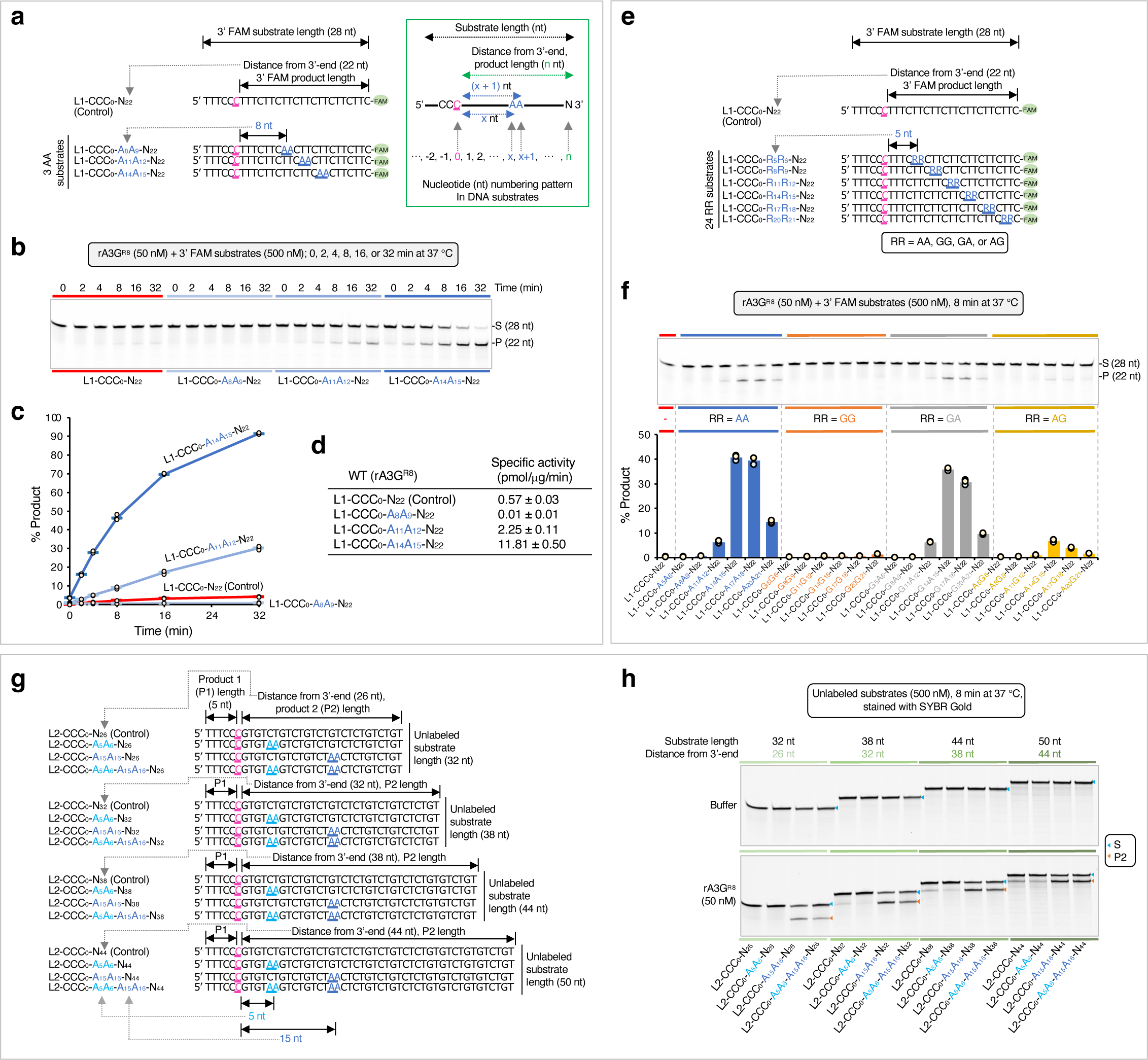
Purine dinucleotide motifs facilitate DNA editing by rA3G. The target-C (in pink) is underlined. Reaction conditions are indicated in the relevant data panels. Each plot shows the average and corresponding data points from three independent experiments. **(a)** Design of a control DNA containing mixed pyrimidine bases with one editing motif CC<u>C</u> placed close to the 5’-end. It is designated as L1-CCC_0_-N_22_ with the target-C at position ‘0’. The subscript ‘22’ specifies the 3’-end nucleotide position relative to the target-C. It also represents the distance (22 nt) between the target-C and the 3’-end, and the length of the editing product. ‘L1’ represents the linear DNA substrate 1 set. Three 28-nt DNA with a single adenine dinucleotide (AA) motif placed in a distance from the target-C are designated as L1-CCC_0_-A_x_A_x+1_-N_22_, where the subscript ‘x’ follows the nucleotide numbering pattern depicted in the inset. **(b)** Gel image and **(c)** Plot of product formation by the four DNA substrates in a time course assay. **(d)** Calculated specific enzyme activities using the linear-range data from the first eight minutes of each reaction. **(e)** Design of 25 3’ 6-FAM labeled 28-nt DNA substrates. RR denotes AA, GG, GA, or AG. **(f)** Gel image and plot of product formation of each DNA substrate in the L1 set. **(g)** Linear DNA substrate 2 (L2) set contains four groups of unlabeled DNA substrates with 32 nt, 38 nt, 44 nt or 50 nt in length. The corresponding distances between the target-C and the 3’-end are 26 nt, 32 nt, 38 nt, and 44 nt. Each group contains four DNA substrates: a control DNA substrate with no AA motifs and three DNA substrates containing individual A_5_A_6_, A_15_A_16_, or a combination of the two AA motifs (A_5_A_6_ and A_15_A_16_). **(h)** Gel images of product formation of each DNA substrate in the L2 set. The quantification data of product formation and specific enzyme activity are presented in **Supplementary Figure 2**.

A linear, 28-nt control DNA in the linear DNA substrate 1 (L1) set was designed with mixed pyrimidine bases (5’-TTTCC<u>C</u>TTTCTTCTTCTTCTTCTTCTTC-FAM 3’, **Fig. 1a**). A single A3G editing motif, CC<u>C,</u> was placed near the 5’-end with a 22-nt distance from 3’-end, reflecting the polarity preference of the A3G deaminase and falling within the ‘dead’ zone^49,50^. Multiple TC motifs were also scattered throughout the sequence. These TC motifs are known to be disfavored by A3G^5,53^ and are unlikely to interfere with editing assays. The nucleotides in the DNA substrate are numbered with the target-C at position ‘0’. Therefore, we designated this control DNA as L1-CCC_0_-N_22_, where the subscript ‘22’ specifies the 3’-end nucleotide position from the target-C (**Fig. 1a** inset). Importantly, this value also represents the distance of the editing motif CC<u>C</u> from the 3’-end and the length of deamination product. A 6-carboxyfluorescein dye (FAM) attached at the 3’ end facilitates the quantification of *in vitro* UDG-dependent deaminase assay.

Three linear AA-containing 28-nt ssDNA in the L1 set were designed with a single AA motif placed at three different locations downstream (in the 3’ direction) of the editing motif CC<u>C</u> (**Fig. 1a**, 1a inset). We designated these AA-containing substrates as L1-CCC_0_-A_8_A_9_-N_22_, L1-CCC_0_-A_11_A_12_-N_22_, and L1-CCC_0_-A_14_A_15_-N_22_, where the numbers following the adenine bases specify the adenine base positions from the target-C.

We utilized a soluble variant of rA3G protein, which was purified from *E. coli*, for the deaminase assay. This variant, documented in our previous studies, is monomeric and largely free of RNA contamination^31,33^. It carries a replacement of N-terminal domain loop 8 (139-CQKRDGPH-146 to 139-AEAG-142, designated as rA3G^R8^) to enhance solubility and has been shown to be catalytically active^31^.

A time course assay was conducted with the control DNA and three AA-containing DNA (**Fig. 1b, 1c**). We observed a significant difference in the editing level among the four DNA substrates. The control DNA L1-CCC_0_-N_22_ has only ∼4% edits. L1-CCC_0_-A_8_A_9_-N_22_ has even lower editing, with ∼0.8% edits. However, L1-CCC_0_-A_14_A_15_-N_22_ has ∼91% edits, followed by L1-CCC_0_-A_11_A_12_-N_22_ with ∼30% edits. Corresponding specific enzyme activities were calculated in the linear product range (**Fig. 1d**), and they varied dramatically from 0.01 pmol/μg/min (L1-CCC_0_-A_8_A_9_-N_22_) to 11.81 pmol/μg/min (L1-CCC_0_-A_14_A_15_-N_22_). The best and the worst rA3G^R8^ specific activity are comparable to those reported for human A3G, about 12 to 15 pmol/μg/min with a 69-nt single stranded DNA, and about 0.07 pmol/μg/min when the editing motif CC<u>C</u> falling within the 30-nt ‘dead’ zone^48,50^.

Next, we extended substrates in the L1 set to include each of the four purine dinucleotide motifs RR (R denotes A or G) and additional RR positions on DNA, while keeping the DNA sequence surrounding the editing motif CC<u>C</u> (5’-TTTCC<u>C</u>TTT) the same in all substrates. Collectively, a panel of 24 ssDNA substrates were derived with six RR positions: R_5_R_6_, R_8_R_9_, R_11_R_12_, R_14_R_15_, R_17_R_18_, and R_20_R_21_ (**Fig. 1e**). Evaluation in the linear product range shows that the top edited substrates were with AA motif, followed by GA motif. The substrates with AG motif also showed low but above-background editing. All GG substrates are poorly edited (**Fig. 1f**). In addition, a pattern of editing efficiency per function of RR position was observed among AA/GA substrates. R_14_A_15_ and R_17_A_18_ are in the best productive positions to promote target-C editing, whereas R_5_A_6_ and R_8_A_9_ are in the non-productive positions.

Following that, we investigated whether increasing distance of the editing motif CC<u>C</u> from 3’-end on longer ssDNA substrates could further facilitate rA3G editing efficiency. We took a low-cost approach and used a panel of unlabeled DNA substrates in combination with a fluorescent SYBR Gold Nucleic Acid Gel Stain detection. Due to relatively weak SYBR Gold signal with pyrimidine only DNA, individual guanine bases (G) were inserted in DNA to boost the staining signal (**Fig. 1g**). Four groups of unlabeled DNA substrates in the linear DNA substrate 2 (L2) set were designed with the distance of CC<u>C</u> from 3’-end increased to 26, 32, 38, or 44 nt. Their substrate lengths were 32, 38, 44, or 50 nt, respectively. Each group contains four substrates including a control DNA without AA motifs and three AA-containing DNA carrying A_5_A_6_ (in the non-productive position), A_15_A_16_ (in the productive position), or both AA motifs (**Fig. 1g, 1h**). The results confirmed that substrates with A_5_A_6_ are poorly edited, whereas substrates with A_15_A_16_ are efficiently edited. The enzyme specific activities are improved to ∼15.14 pmol/μg/min with L2-CCC_0_-A_15_A_16_-N_32_, and then it stays close to this value as the distance of the editing motif CC<u>C</u> from 3’-end increases (such as ∼14.31 pmol/μg/min with L2-CCC_0_-A_15_A_16_-N_44_, **Supplementary Fig. 2**). Additionally, two AA motifs, A_5_A_6_ and A_15_A_16_, simultaneously generate a combined effect on one CC<u>C</u> motif.

We also observed that, as the distance of the editing motif CC<u>C</u> from 3’-end increases, a substantial number of edits are generated even in the control DNA that contains no AA motifs (such as L2-CCC_0_-N_44_ with the specific enzyme activity of 6.59 pmol/μg/min, **Supplementary Fig. 2**). Consequently, AA-facilitated editing is less pronounced, with the specific enzyme activity being about two-fold higher (14.31 pmol/μg/min, **Supplementary Fig. 2**). These results suggest that with increasing distance of CC<u>C</u> from 3’-end, AA-independent interactions between rA3G and substrate DNA also increase, leading to efficient DNA capture and target-C deamination in the absence of AA dinucleotide motifs.

In summary, we find that a single AA or GA motif can facilitate rA3G^R8^ editing efficiency on its target-C. AA/GA-facilitated editing is dictated by their position from the target-C with R_14_A_15_ to R_17_A_18_ in the best productive positions (specific enzyme activities ∼11.8 to ∼15.14 pmol/μg/min), and with R_5_A_6_ to R_8_A_9_ in the non-productive (or inhibitory) positions (specific enzyme activities ∼0.01 to 0.41 pmol/μg/min). The magnitude of AA-facilitated editing is also influenced by the distance of the editing motif CC<u>C</u> to the 3’-end. As this distance increases, AA-facilitated editing is attenuated, while AA-independent editing is boosted. Lastly, two adjacent AA motifs can generate a combined effect on a single CC<u>C</u> motif.

### Overall structures of rA3G bound with ssDNA containing AA or GA motif

Prior to crystallization trials, we determined the minimal productive AA position of deamination on a target-C. We compared a panel of 17 substrates in the linear DNA substrate 3 (L3) set, each carrying a single AA motif placed from 5 nt to 21 nt downstream of the editing motif CC<u>C</u> (**Fig. 2a**). The results show that 10 nt (A_10_A_11_) is the minimal distance to elicit AA-facilitated editing function (**Fig. 2b**). With this information, crystallization trials were carried out using the catalytically inactive rA3G^R8/E259A^ and ssDNA with AA or GA positioned at R_10_A_11_, R_11_A_12_, or R_14_A_15_. The lengths of the DNA substrate were shortened by removing the last four nucleotides at the 3’-end (**Fig. 2c, 2d**). Additionally, GA-containing DNA sequences were further modified to replace guanine bases outside of the GA motif with thymine bases (**Fig. 2d**). The best diffracting crystals were obtained with A_10_A_11_- and G_11_A_12_-containing DNA (**Fig. 2c, 2d**), and their structures were determined.

**Figure 2.**
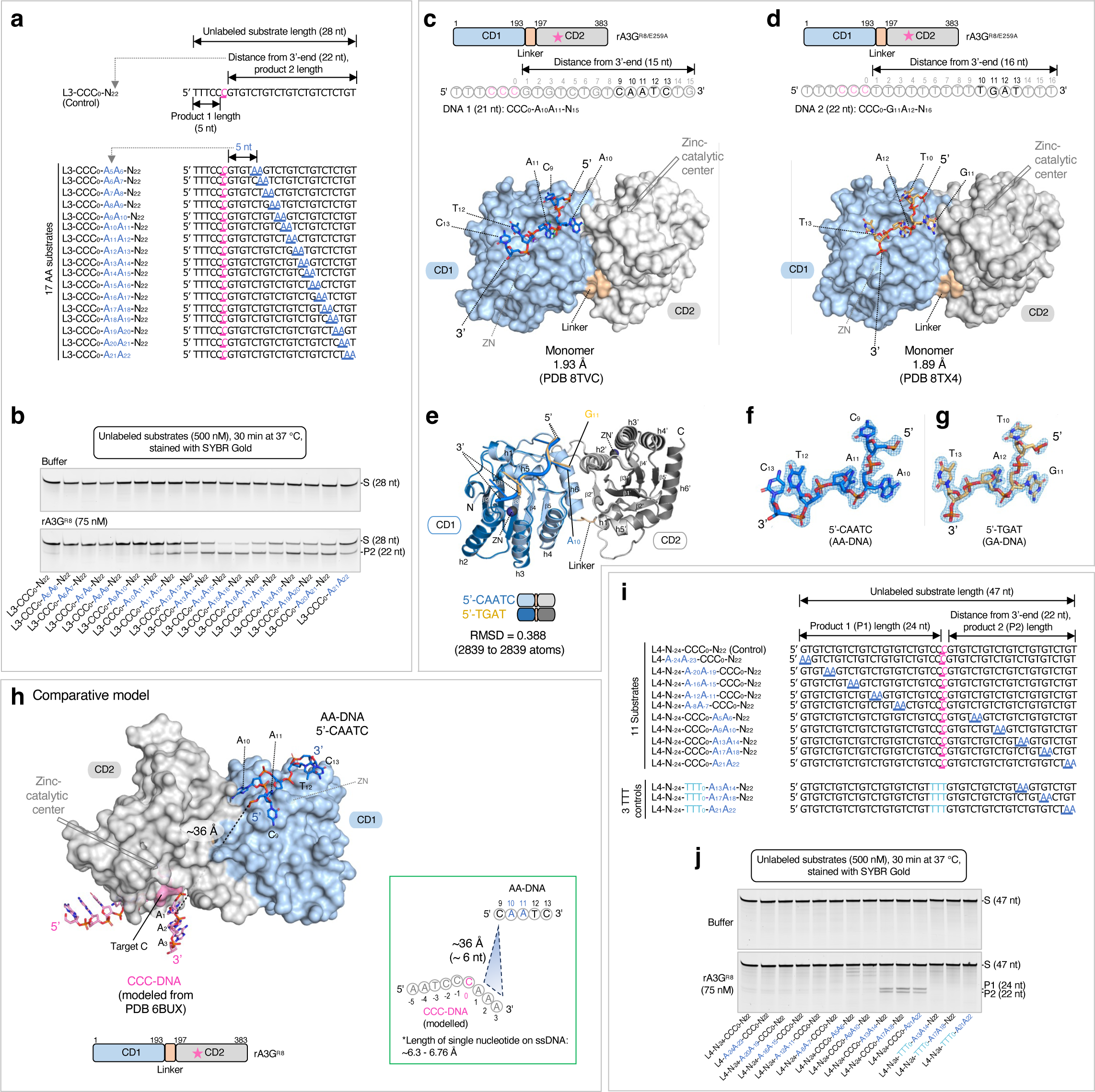
Crystal structures of rA3G^R8/E259A^ in complex with short DNA sequence 5’-CAATC (AA-DNA) or 5’-TGAT (GA-DNA). **(a)** Linear DNA substrate 3 (L3) set contains 17 unlabeled 28-nt DNA substrates with a single AA motif placed stepwise downstream from the target-C. A control DNA with no AA motif (L3-CCC_0_-N_22_) is also shown. **(b)** Gel images of product formation, showing that the minimal productive AA position is A_10_A_11_. **(c) (d)** Surface and stick representation of the structure of rA3G^R8/E259A^ in complex with a short DNA sequence 5’-CAATC (in marine sticks) or 5’-TGAT (in light orange sticks). Zinc-catalytic residue E259A is marked by a pink star in the schematic diagram. Nucleotides in black are resolved. **(e)** Superimposition of the two models showing they are essentially the same with a subtle difference between the nucleotide A_10_ in 5’-C_9_A_10_A_11_T_12_C_13_ and G_11_ in 5’-T_10_G_11_A_12_T_13_. **(f)** (**g)** 2Fo-Fc electron density map of the resolved short DNA sequence 5’-CAATC or 5’-TGAT contoured at 1.5 σ level. **(h)** Comparative modeling of rA3G bound to AA-DNA (this study) and the editing motif CCC-DNA (modeled from PDB 6BUX^44^). The straight-line distance between A_1_ in the CC<u>C</u>-DNA (in pink sticks, modeled from PDB 6BUX^44^ and C_9_ in the AA-DNA (in marine sticks) is indicated by a black dotted line and shown in the inset. Length per nucleotide ranging from 6.3 - 6.76 Å^54,55^ is used to convert distance to number of nucleotides. **(i)** Linear DNA substrate 4 (L4) set contains 10 unlabeled 47-nt DNA substrates with a single AA motif placed systematically upstream and downstream from the target-C. A control DNA with no AA motif is also shown. In the three TTT controls, the editing motif CCC was replaced by TTT. **(j)** Gel images of product formation, showing that the AA-facilitated editing occurs only in positions A_13_A_14_, A_17_A_18_, and A_21_A_22_ downstream of the editing motif CCC.

Both structures of the rA3G^R8/E259A^-DNA complexes are monomer complexes with one rA3G^R8/E259A^ molecule bound to one DNA molecule, with a resolution determined at 1.93 Å and 1.89 Å, respectively (**Fig. 2c, 2d**). Five nucleotides spanning over the AA motif (5’-C_9_A_10_A_11_T_12_C_13_) and four nucleotides over the GA motif (5’-T_10_G_11_A_12_T_13_) were built into the electron density unambiguously (**Fig. 2f, 2g**). However, the editing motif CC<u>C</u> and the rest of the DNA at the 5’-end remained unresolved in both structures, likely due to their flexibility because of no strong binding of the CC<u>C</u> motif to the CD2 domain. Superimposition of the two complex structures yielded rmsd of 0.388 Å (2839 to 2839 atoms, **Fig. 2e**), indicating they are essentially the same structure. A subtle but noticeable difference is seen between A_10_ of the AA motif (A_10_A_11_) and G_11_ of the GA motif (G_11_A_12_) (**Fig. 2e, Supplementary Fig. 3**), which is expected for the guanine base G_11_ of the GA motif to fit into the groove that tightly binds the adenine base A_10_ of the AA motif. The structure of rA3G^R8/E259A^ bound to the AA-containing DNA (resolution 1. 93 Å) is used to describe the protein-DNA interactions in the following sections.

To place the editing motif CCC into the context of the full-length A3G structure, we carried out comparative modeling based on our rA3G bound to AA-DNA structure, alongside a previously determined structure of human A3G-CD2 bound to the editing motif CC<u>C</u> (referred to as ‘CCC-DNA’, modelled from PDB 6BUX^44^). The comparative model predicted that the AA-DNA fragment (predominately bound by rA3G-CD1) positions its 5’-end (C9, **Fig. 2h**) in the general direction of the 3’-end of the editing motif CC<u>C</u>, as modeled with rA3G-CD2 (**Fig. 2h**). This prediction is consistent with the inherent directionality of CC<u>C</u> and AA motifs in our DNA substrates (**Fig. 1e, 1g**). To further validate the polar arrangement, we utilized a panel of 11 unlabeled 47-nt DNA substrates in the linear DNA substrate 4 (L4) set with a single AA motif systematically positioned upstream or downstream of the editing motif CC<u>C</u> (**Fig. 2i**). Our observations show that AA-facilitated editing occurs only in positions A_13_A_14_, A_17_A_18_, and A_21_A_22_ downstream of the editing motif CC<u>C</u> (**Fig. 2j**), aligning with the spatial organization predicted in the comparative model.

We further estimated the distance between the editing motif CC<u>C</u> and the AA motif by measuring the straight-line distance between A_1_ in CCC-DNA and C_9_ in AA-DNA (**Fig. 2h** and its inset), and determined it to be ∼36 Å. Using a nucleotide length of 6.3 or 6.76 Å for ssDNA^54,55^, this distance corresponds to roughly 6 nt between A_1_ and C_9_. In a real situation, the distance is expected to be longer than 6 nt as it should not follow a straight line connecting A_1_ and C_9_ due to the protein surface features. Therefore, this estimation aligns well with our experimentally determined minimal distance of 7 nt between A_1_ and C_9_ for the productive configuration, which is equivalent to A_10_A_11_ (**Fig. 2b**).

### Detailed interactions between rA3G and DNA

In the co-crystal structure of rA3G^R8/E259A^ bound to AA-containing DNA, the short 5-nt DNA centered around the AA dinucleotide (5’-C_9_A_10_A_11_T_12_C_13_) out of the 21-nt ssDNA are clearly visible (**Fig. 2f**). The AA dinucleotide bases (A_10_ and A_11_) are inserted deep inside the protein, the nucleotides before and after A_10_A_11_ (i.e. C_9_, T_12_ and C_13_) are bound on the protein surface. The rA3G binding interface for the 5-nt DNA is composed of 15 amino acid residues, 13 residues of which are located on the CD1 loops near CD1 Zn-center (loops 1, 3, 5, and 7), with the remaining 2 residues coming from CD2 (**Fig. 3a-3e**). A hydrophobic groove conformed between CD1 and CD2 binds to the 5’-A (A_10_), and a hydrophobic cave-like pocket on CD1 binds to the 3’-A (A_11_) of the AA dinucleotide. The groove donates five residues (I26, F126, W127 on CD1, and F268 and K270 on CD2) to interact with A_10_ through mostly hydrophobic interactions and only one hydrogen bond (**Fig. 3b**). The cave-like pocket of CD1 interacts with A_11_ via hydrophobic packing and four strong hydrogen bonds through eleven CD1 residues, including 25-PILS-28 on loop 1, Y59 on loop 3, W94 on loop 5, and 123-LYYFW-127 on loop 7 (**Fig. 3c, 3e**). Additionally, five CD1 residues, 24-RPILS-28 (loop 1), form a small surface area that interacts with C_9_ (**Fig. 3d**). Two CD1 residues, 59-YP-60 (CD1 loop 3), have weak interactions with T_12_C_13_ (**Fig. 3e**).

**Figure 3.**
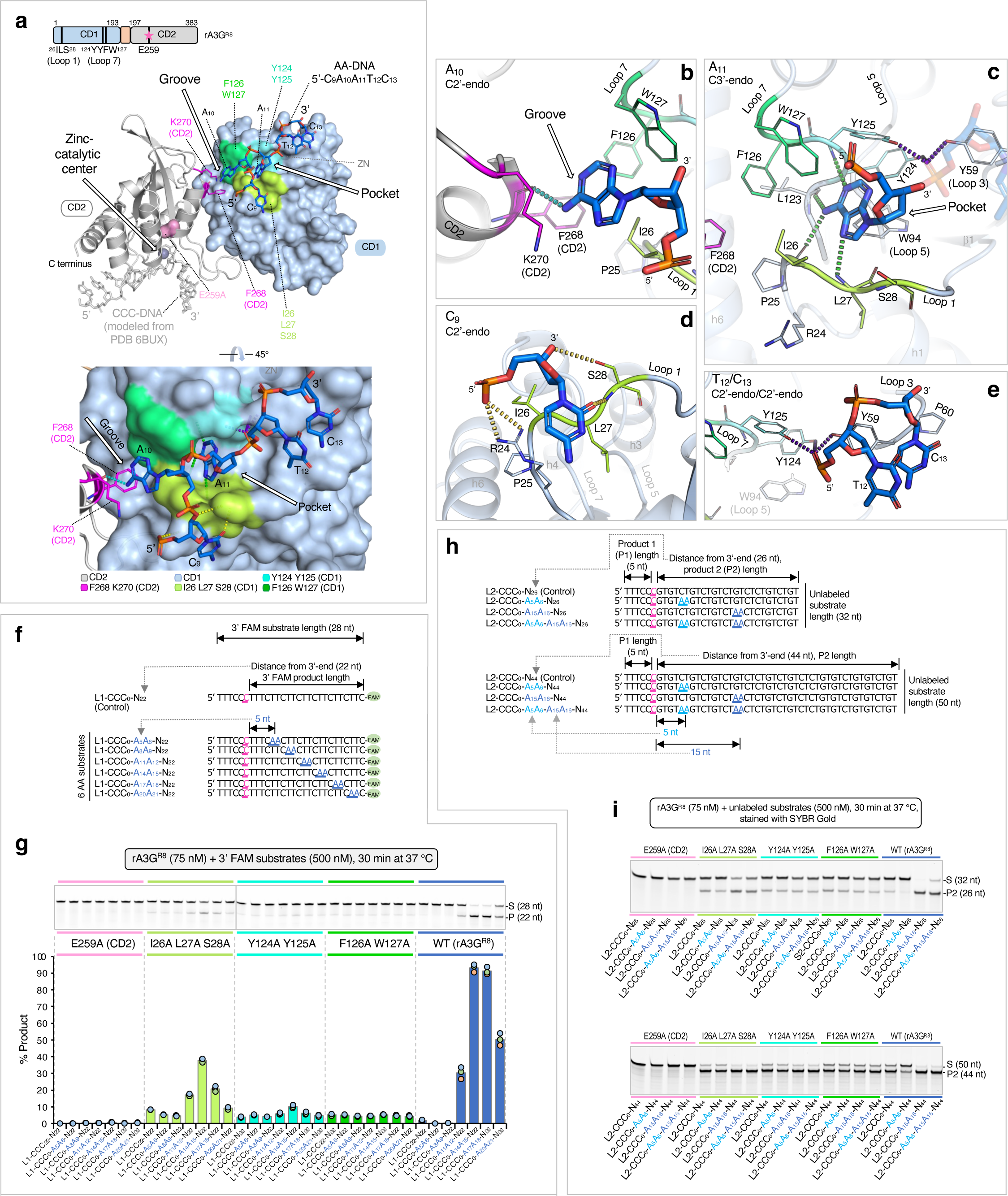
Interactions between rA3G and AA-DNA. **(a)** Surface and ribbon representation of the CD1 domain (in light blue surface) and the CD2 domain (in gray ribbon) bound to AA-DNA 5’-C_9_A_10_A_11_T_12_C_13_ (in marine sticks). CCC-DNA (in light gray sticks) is modeled from PDB 6BUX^44^. Three surface patches surrounding the A_10_A_11_ binding pocket/groove are formed by multiple amino acid residues (^26^ILS^28, 124^YY^125^, and ^126^FW^127^). A catalytically inactive mutation E259A (in pink surface) is shown on the CD2 domain structure (in gray ribbon). Two CD2 residues F268 and K270 are shown (in magenta sticks). **(b)**, **(c)**, **(d)**, **(e)** Ribbon and stick representations of detailed interactions between rA3G and DNA. The color theme is the same as in (a). Hydrogen bonds are indicated by dashed lines. **(f)** Sequences of the L1 DNA substrates containing a single AA motif and a control DNA containing no AA motif. **(g)** Gel image and plot of product formation, showing much reduced AA-facilitated editing by three mutants. **(h)** Two groups of L2 DNA substrates with 32-nt and 50-nt in length, respectively. Each group contains a control DNA substrate with no AA motifs and three DNA substrates containing individual A5A6, A15A16, or a combination of the two AA motifs (A5A6 and A15A16). **(i)** Gel images of product formation with 32-nt and 50-nt DNA. _3_

Overall, if comparing the rA3G structures in complex with the AA-DNA (5’-C_9_A_10_A_11_T_12_C_13_) vs rArA-RNA (5’-rU_4_rA_5_rA_6_rU_7_rU_8_)^33^, the first four DNA nucleotides (C_9_A_10_A_11_T_12_) align very well with the four RNA nucleotides (rU_4_rA_5_rA_6_rU_7_) and show nearly identical interactions with rA3G, with the fifth DNA nucleotide C_13_ adopting different interactions with rA3G from the corresponding RNA nucleotide rU_8_ (**Supplementary Fig. 4**a). These results indicate that rA3G binds to the AA dinucleotide and the immediate 5’ and 3’-side nucleotides very similarly for both DNA and RNA. The only noticeable differences are that RNA forms hydrogen bonds with rA3G S28 sidechain and main-chain N and rA3G G29 main-chain N through the 2’-OH of the sugar moiety of rU_4_ and rA_6_ (yellow sticks in **Supplementary Fig. 4**b), which are absent in DNA due to the lack of 2’-OH. For the fifth DNA nucleotide C_13_, it turns in a different direction from its equivalent RNA nucleotide rU_8_ in such a way that C_13_ packs with T_12_. Such packing interaction between C_13_ and T_12_ should not allowed in RNA due to the presence of 2’-OH of rU_7_ that would clash with rU_8_.

To verify the importance of AA binding residues in AA-facilitated editing efficiency, we generated alanine mutations on seven CD1 amino acid residues that engage with the AA dinucleotide of the bound DNA, including I26A/L27A/S28A, Y124A/Y125A, and F126A/W127A. A wild type (rA3G^R8^) and a catalytically inactive mutant (rA3G^R8/E259A^) were used as positive and negative controls (**Supplementary Fig. 5**a). Using a panel of 3’ FAM labeled AA-containing substrates (in the L1 set), the results show that while rA3G^R8/E259A^ is catalytically dead, two of the three CD1 mutants, Y124A/Y125A and F126A/W127A, displayed only basal level editing function (**Fig. 3f, 3g**), indicating the critical importance of AA binding residues 124-YYFW-127 of CD1 in facilitating efficient editing by CD2. However, AA-facilitated editing is only partially lost in the mutant I26A/L27A/S28A, suggesting that these residues are less critical, and the mutant may still retain partial binding to AA.

Further validation was carried out using two groups of long DNA substrates (in the L2 set) with increased distances between the editing motif CC<u>C</u> and the 3’-end (26 nt and 44 nt, **Fig. 3h**). Similar results were obtained that theses mutants show defective in AA-facilitated editing (**Fig. 3i**). However, significant number of edits are generated by these mutants in the long DNA substrates that have 44 nt between the editing motif CC<u>C</u> and the 3’-end, indicating that AA-independent editing is largely unaffected in these mutants.

We also generated alanine mutations on CD2 residues F268 and K270 that participate in binding to the adenine base A_10_ (**Fig. 4a**, **Supplementary Fig. 5**b) and tested it with a panel of 3’ FAM labeled RR-containing substrates (**Fig. 4b**). Comparing to the wild type, the mutant F268A/K270A has an overall reduced editing efficiency when RR is in the productive positions (R_11_R_12_, R_14_R_15_, R_17_R_18_, and R_20_R_21_, **Fig. 4c, 4d**), indicating that the impairment in AA-binding leads to an impaired AA-facilitated editing. Interestingly, it has an overall slightly increased editing efficiency when RR is in the non-productive positions (R_5_R_6_ and R_8_R_9_, **Fig. 4c, 4d**). Further examination of the minimal AA register using eight DNA substrates in the L3 substrate series (**Fig. 4e, 4f**) shows that the minimal AA register for productive DNA editing is shortened to ∼6 nt (A_6_A_7_). These observations suggest that the physical barrier between the CCC motif and AA motif has changed, possibly due to weakened interface rigidity between the CD1-CD2 domains. This allows CD2 greater freedom to rotate relative to CD1, enabling it to interact with DNA more flexibly and reach the CC<u>C</u> motif at a shorter distance.

**Figure 4.**
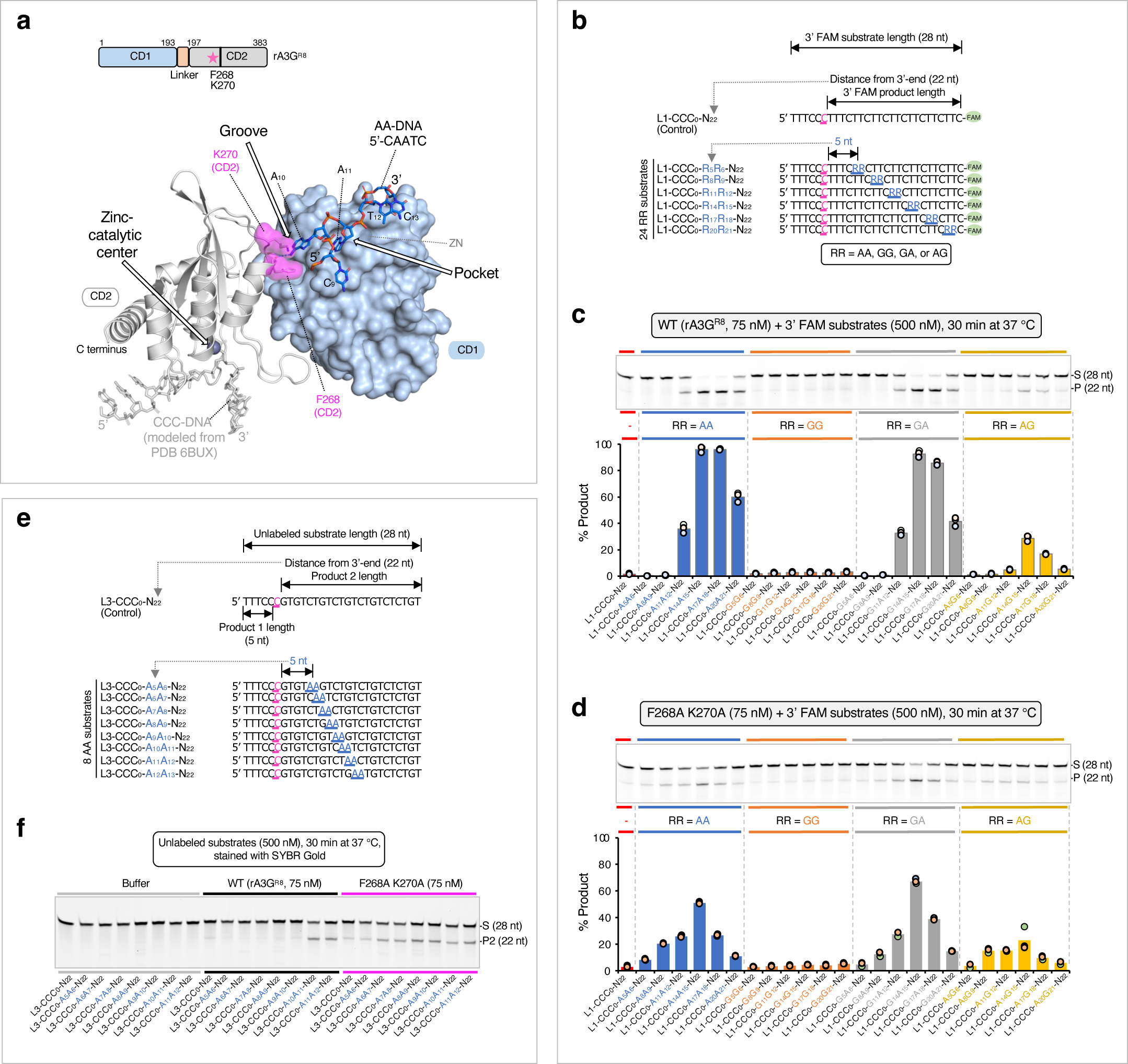
AA/GA-facilitated DNA editing performed with rA3G^R8^ carrying F268A and K270A mutations. **(a)** Surface and ribbon representation of the CD1 domain (in light blue surface) and the CD2 domain (in gray ribbon) bound to AA-DNA 5’-CAATC (in marine sticks). CCC-DNA (in light gray sticks) is modeled from PDB 6BUX^44^. Two CD2 residues F268 and K270 are shown (in magenta surface). **(b)** Sequences of the L1 DNA substrates, where RR denotes AA, GG, GA, or AG. **(c)** Gel image and plot of product formation by the wild-type rA3G^R8^. **(d)** Gel image and plot of product formation by the mutant rA3G^R8/F268A/K270A^. **(e)** Sequences of the first 8 DNA substrates in the L3 DNA substrate set. **(f)** Gel images of product formation showing the minimal productive AA position is A_10_A_11_ for the wild-type rA3G^R^^8^ and A_6_A_7_ for the F268A K270A mutant.

### Editing property of a hyperactive rA3G variant

We investigated whether rA3G carrying a hyperactive catalytic domain could override or escape from the AA-facilitated editing. We constructed a rA3G^R8^ variant carrying two mutations on its CD2 domain, P247K and Q317K (**Fig. 5a, 5b**). Their corresponding mutations P247K and Q318K from human A3G have shown to contribute to the hyperactivity of the human A3G CD2 catalytic domain^44^. The rA3G^R8/P247K/Q317K^ variant displays much enhanced editing efficiency on an AA-containing DNA substrate L1-CCC_0_-A_14_A_15_-N_22_ (**Fig. 5c, 5d, 5g**). Its enzyme specific activity reached 101.82 pmol/μg/min, about 8.6-fold higher than that of the wild-type rA3G^R8^. Dramatic increase in editing efficiency was also observed in other AA-containing DNA and in the control DNA (**Fig. 5g**). Despite the overall enhanced efficiency, the substrate rank order remained the same as that of the wild type: L1-CCC_0_-A_14_A_15_-N_22_ is still the best substrate, followed by L1-CCC_0_-A_11_A_12_-N_22_, control DNA, and L1-CCC_0_-A_8_A_9_-N_22_. Using the complete panel of 25 FAM labeled DNA (**Fig. 5e, 5f**), we show that the hyperactive rA3G variant displays a similar pattern as that of the wild type, albeit under a much lower enzyme concentration. GG-containing substrates remained to be poor substrates. Of note, the editing efficiency on AG substrates were disproportionally enhanced.

**Figure 5.**
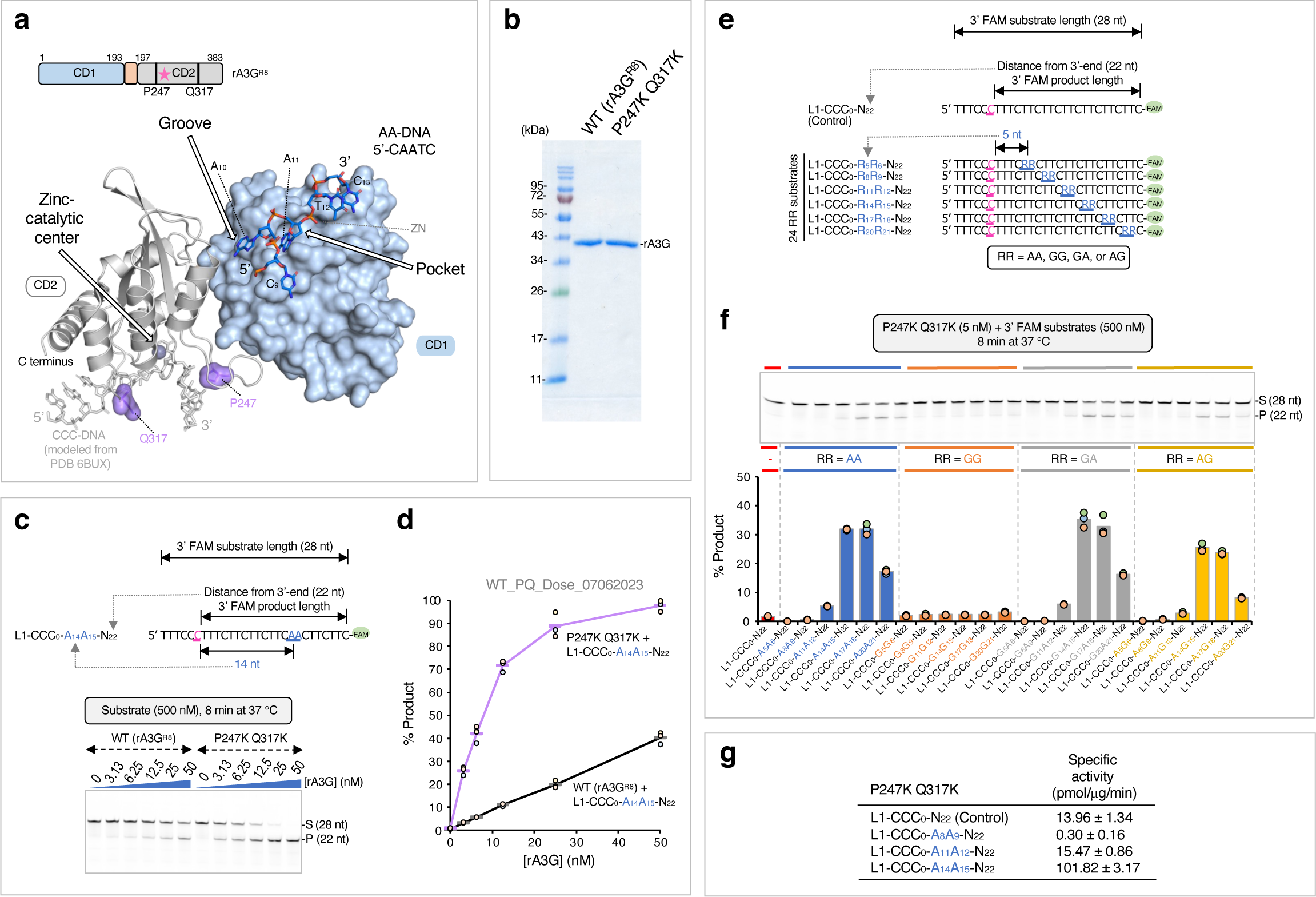
AA/GA-facilitated DNA editing performed with rA3G^R8^ carrying P247K and Q317K mutations. **(a)** Surface and ribbon representation of the CD1 domain (in light blue surface) and the CD2 domain (in gray ribbon) bound to AA-DNA 5’-CAATC (in marine sticks). CCC-DNA (in light gray sticks) is modeled from PDB 6BUX^44^. Two previously characterized residues P247 and Q317 (in purple surface) are shown on the CD2 domain (in gray ribbon). Corresponding mutations P247K and Q318K cause a hyperactive phenotype in human A3G CD2 domain^44^. **(b)** SDS-PAGE gel image showing the purified rA3G^R8/ P247K/Q317K^ and the wild-type rA3G^R8^. **(c)** Gel image and **(d)** plot of product formation in a dose response assay, showing enhanced editing by rA3G^R^^8/P247K/Q317K^. The 3’ FAM labeled DNA substrate sequence, L1-CCC_0_-A_14_A_15_-N_22_, is also shown. **(e)** Sequences of the L1 DNA substrates, where RR denotes AA, GG, GA, or AG. **(f)** Gel image and plot of product formation, showing that rA3G^R8/P248K/Q317K^ retains the AA/GA-facilitated editing pattern and displays an enhanced AG-facilitated editing pattern. **g)** Calculated specific enzyme activity of four 3’ FAM labeled DNA substrates by rA3G^R8/P248K/Q317K^.

### RNA inhibition of AA-facilitated DNA editing

As shown earlier that rA3G uses the same structural elements to bind AA (and GA) in DNA and RNA in a similar fashion (**Fig. 6a**, **Supplementary Fig. 4**), and that the AA binding by A3G dictates the editing efficiency of the target-C located upstream of the AA dinucleotide. This provides a plausible structural explanation to prior data on RNA inhibition of A3G DNA editing function^56–60^. To verify this, we tested a panel of AA-DNA substrates with six AA positions (A_5_A_6_, A_8_A_9_, A_11_A_12_, A_14_A_15_, A_17_A_18_, and A_20_A_21_), and two 10-nt RNA competitors, rArA-RNA, and rUrU-RNA (5’-rUrUrUrUrArArUrUrUrU and 5’-rUrUrUrUrUrUrUrUrUrU, respectively). We used the hyperactive rA3G^R8/P247K/Q317K^ variant to perform the competition assay (**Fig. 6b, 6c**). The results show that rArA-RNA (but not rUrU-RNA) inhibits AA-facilitated DNA editing. We further tested RNA inhibition of AA-facilitated DNA editing using one of the DNA substrates, L1-CCC_0_-A_17_A_18_-N_22_, and a set of six 10-nt RNA competitors (5’-rUrUrUrUrNrNrUrUrUrU, where rNrN denotes rArA, rUrA, rUrU, rGrG, rGrA, or rArG. We find that rArA or rGrA-containing RNA competitors cause substantial reduction in the AA-facilitated DNA editing, which supports that rArA/rGrA RNA compete with AA-DNA for the same binding site on rA3G. Because many cellular RNAs contain unpaired rArA motif, the cellular rArA-containing RNA bound to A3G-CD1 is expected to inhibit the editing activity of A3G if they are not displaced.

**Figure 6.**
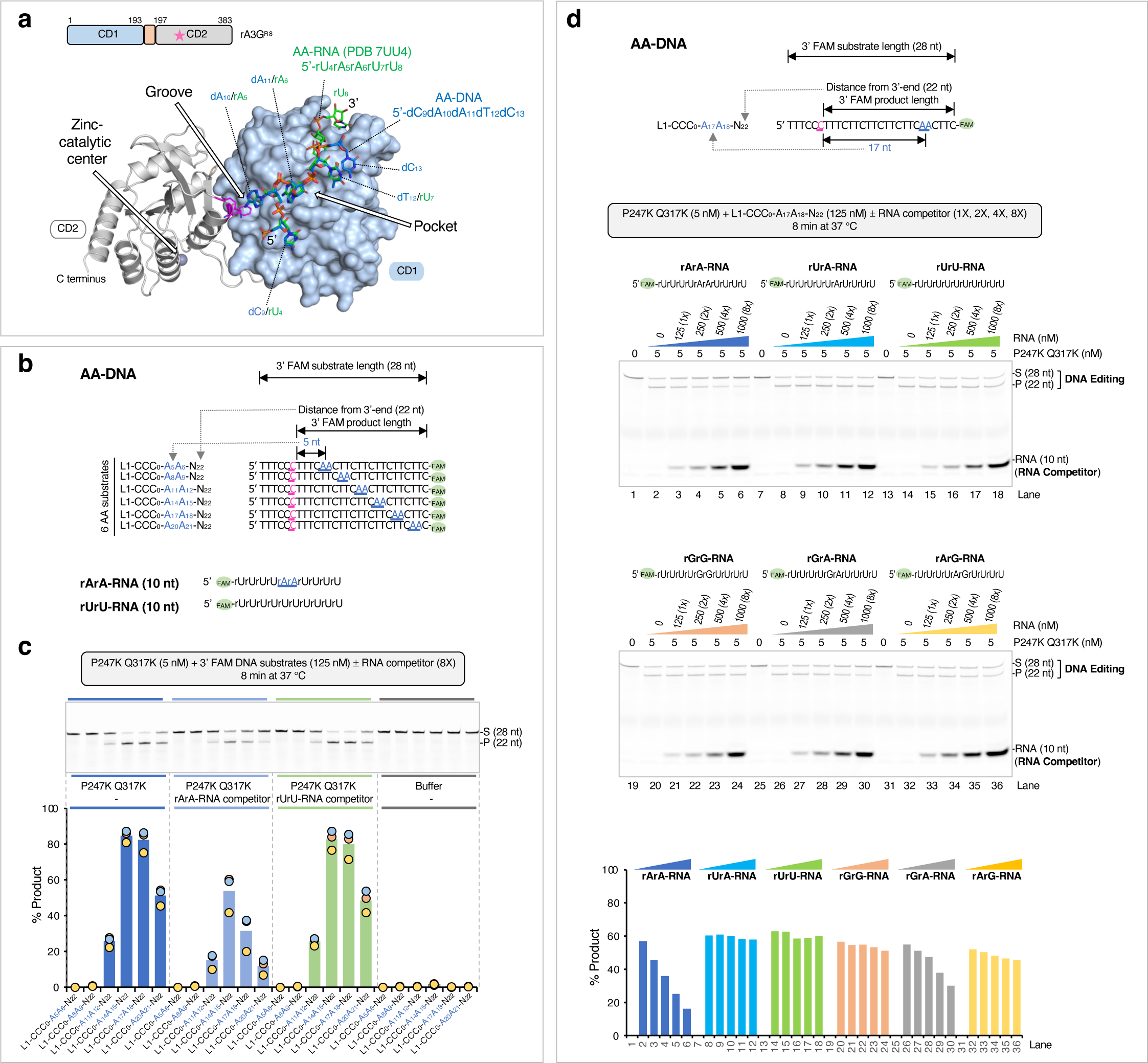
Inhibition of AA-facilitated DNA editing by RNA. DNA and RNA were mixed at the indicated molar ratio in the reaction mixture, and the assays were performed using the hyperactive variant rA3G^R8/P247K/Q317K^. **(a)** Overlapping binding surfaces for the AA motif on DNA and the rArA motif on RNA with rmsd 0.193 (2660 atom pairs). See **Supplementary Fig. 4** for further details. **(b)** Sequences of six L1-AA DNA substrates and two 10-nt RNA competitors used in the inhibition assay. **(c)** Gel image and plot of product formation, showing reduced editing in the presence of rArA RNA and no effect in the presence of rUrU RNA. **(d)** Inhibition assay with six 10-nt RNA competitors. rArA RNA and rGrA RNA show inhibitory effects on AA DNA (L1-CCC_0_-A_17_A_18_-N_22_).

### AA-facilitated DNA editing in hairpin forming sequences

A3A and A3B have been shown to edit the target-C presented in both linear and short hairpin loop DNA, displaying a preference for the hairpin loop substrates. However, there is no report showing that A3G can edit the CC<u>C</u> motif in a hairpin loop DNA structure. Furthermore, A3G-DNA structures with linear DNA show no base-paring, as seen in A3A/DNA structures^44,61,62^. On the other hand, it has been reported that A3G can edit the target-C in the hairpin loop of an RNA hairpin^27–30^.

From comparative modeling of our rA3G bound to AA-DNA structure and a previously determined structure of human APOBEC3A bound to a tetraloop DNA hairpin (PDB 8FIK)^62^, we hypothesized that a DNA hairpin substrate with a long stem-length and a 3’overhang (for presenting AA motifs to the non-catalytic CD1 domain) could potentially be edited by rA3G (**Fig. 7a**). We used the hyperactive rA3G^R8/P247K/Q317K^ variant to test this hypothesis.

**Figure 7.**
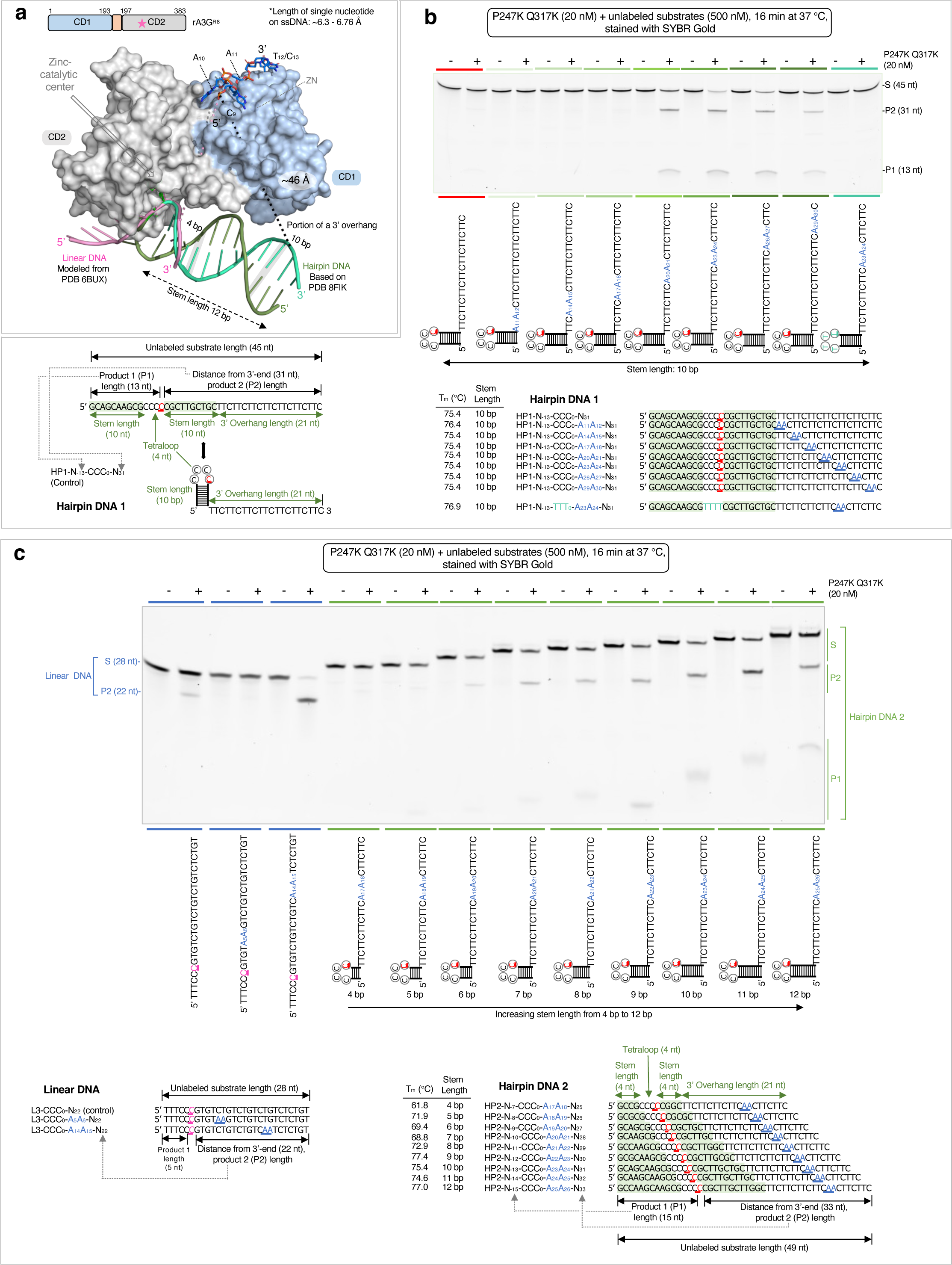
AA-facilitated DNA editing in hairpin forming sequences performed with the hyperactive variant rA3G^R8/P247K/Q317K^. **(a)** Comparative modeling of rA3G bound to AA-DNA (this study), the editing motif CCC-DNA (in pink, modeled from PDB 6BUX^44^) and a DNA hairpin structure with a tetraloop and a 12-bp stem length (in green, based on PDB 8FIK^62^). A potential connection path (∼46 Å) between the hairpin DNA (with a 10-bp stem length) and the AA motif is indicated by a black dotted line and labeled as ‘portion of a 3’overhang’. **(b)** Location of the AA motif on 3’ overhang affects editing efficiency. Deamination activity was monitored on hairpin DNA 1 substrates carrying a 10-bp hairpin stem, a tetraloop CCCC, and a 3’overhang. A single AA motif was placed at various locations on the 3’ overhang. A negative control with 5’-TTTT in the hairpin loop and a single A_23_A_24_ motif in the 3’overhang was included. The results show that the top three edited hairpin DNA with the stem length of 10 bp are those carrying a single AA motif A_20_A_21_, A_23_A_24_, or A_26_A_27_. **(c)** Effect of stem length on editing efficiency. Deamination activity was monitored on hairpin DNA 2 substrates with varying stem lengths from 4 bp to 12 bp. The position of the AA motif was kept the same in all substrates as the position of A_23_A_24_ in the 10-bp stem loop DNA substrate. Three linear DNA substrates were included as linear DNA controls. The results show that hairpin DNA with short stem lengths (4 bp or 5 bp) are poorly edited, whereas hairpin DNA with long stem lengths (10 bp or 11 bp) are well edited.

We designed a control DNA substrate (hairpin DNA substrate 1 or HP1) with the editing motif CC<u>C</u> situated in the loop region of a hairpin structure with a 10-bp hairpin stem length, 5’-GCAGCAAGCG(CCC<u>C</u>) CGCTTGCTGC. The hairpin DNA also carries a 21-nt 3’overhang to boost its interaction with CD1 (**Fig. 7b**). Seven AA-containing DNA were designed with the AA motifs placed at various locations in the 3’overhang.

All annealed hairpin DNA were essentially monomeric (**Supplementary Fig. 6**a) with the estimated T_m_ between 75.4 to 76.9 °C (**Fig. 7b**). The results show that the hairpin DNA are not efficiently edited without AA motif or with AA motif located from 11-12 nt to 17-18 nt downstream from the target C (substrates with A_11_A_12_, A_14_A_15_, or A_17_A_18_). Instead, efficient editing was only observed when the AA motif is positioned at a distance longer than 20-21 nt between A_20_A_21_ and A_29_A_30_.

Comparing with linear DNA substrates, the minimal AA register for the productive DNA editing has changed from A_10_A_11_ to be longer than A_17_A_18_. This is likely caused by the difference between the rigid form of the hairpin duplex and the flexible linear DNA, as well as their spatial arrangement with the rA3G protein (**Fig. 7a, Supplementary Fig. 9**). In the comparative model, the straight-line distance between the 3’-end of the hairpin stem and the 5’-end of the AA-DNA is about ∼46 Å, which is equivalent to 7 or 8 nt. This model shows that A_17_A_18_ does not have sufficient linear space between A_17_A_18_ bound at CD1 and the target CC<u>C</u> in the stem-loop bound at CD2 active site. It requires at least A_20_A_21_ to cover the distance and support the AA-facilitated editing of the target-C in this hairpin DNA. We further tested the effect of stem-length on editing efficiency. Nine unlabeled hairpin DNA substrates were designed to carry varies stem-lengths from 4 bp to 12 bp (hairpin DNA 2 or HP2 set, **Fig. 7c**).

Additionally, they all have the same 3’overhang sequence with one AA motif placed at the 13 −14 nt position from the 3’-end of the hairpin stem. All annealed hairpin DNA of different hairpin stem lengths were monomeric with the estimated T_m_ between 61.8 to 77.4 °C (**Fig, 7c**, **Supplementary Fig. 6**b). Three unlabeled linear DNA from the L3 set, L3-CCC_0_-N_22_, L3-CCC_0_-A_5_A_6_, and L3-CCC_0_-A_14_A_15_-N_22_ (**Fig. 2a, 7c**) were also included as the linear DNA controls. Our results show that the linear DNA displayed the expected editing pattern with near complete editing in L3-CCC_0_-A_14_A_15_-N_22_ and very little editing in L3-CCC_0_-A_5_A_6_-N_22_. Nine hairpin DNA2 substrates also displayed a dramatic difference in editing efficiency. Hairpin DNA with short stem lengths are poorly edited (4 bp and 5 bp), whereas hairpin DNA with long stem lengths are efficiently edited with the top hairpin DNA carrying 11 bp, followed by 10 bp and 9 bp. DNA with the longest hairpin stem tested (12 bp) was less well edited.

Additional tests were carried out with hairpin DNA substrates containing a fixed hairpin stem-length of 4 bp (HP3 set) or 11 bp (HP4 set), with varying AA positions in the 3’overhang (**Supplementary Fig. 7, 8**).

The results show that hairpin DNA with the 4-bp stem length are overall poor substrates, whereas hairpin DNA with the 11-bp stem length yield similar results to the 10-bp stem length (**Fig. 6b**). These findings align with our model predictions (**Supplementary Fig. 9**), which suggest that proper stem lengths upstream of the AA motif enable the CCC-loop at the end of the stem to reach CD2 for deamination. Consequently, the principle governing the spatial requirement between the AA motif recognized largely by CD1 and the target cytosine edited by CD2’s active site is consistent across both linear DNA and hairpin DNA, despite differences in the number of nucleotides between the AA motif and the target cytosine.

## Discussion

In this study, we have demonstrated that rA3G editing of target-C in both linear and hairpin loop sequences is significantly influenced by the presence of AA and GA dinucleotides at certain distance downstream (but not upstream) of the target-C. We have also provided the mechanistic understanding for these biochemical observations through determination of two co-crystal structures of the full-length rA3G in complex with AA- or GA-containing ssDNA sequences. These structures reveal how A3G uses its non-catalytic CD1 domain to capture the substrate through recognition of the AA/GA motifs with a specific orientation to present the target-C to the distally located active site on the catalytic CD2 domain for deamination.

The AA/GA-facilitated DNA editing is directional and dynamic, supporting previous findings regarding the directionality and processivity features of A3G catalysis on ssDNA. Multiple AA, GA motifs present in the DNA substrates in previous reports would allow A3G to bind at multiple locations on these DNA substrates, thus rationalizes A3G “promiscuous” DNA binding property. The biochemical and structural data described here, together with the prior information reported from other studies^44,48,50,52^, support a mechanism that CD1 scans, recognizes, and captures the AA/GA motif in ssDNA region, which allow CD2 sufficient time to hit on those CC<u>C</u> motifs located 5’-upstream within the editing window to catalyze the deamination on the target-Cs (**Fig. 8**). DNA secondary structure can affect the geometrical distance of AA/GA and CC<u>C</u> on DNA, which in turn, affects the register of the productive distance between AA/GA and the target-C (**Supplement Fig. 9**).

**Figure 8.**
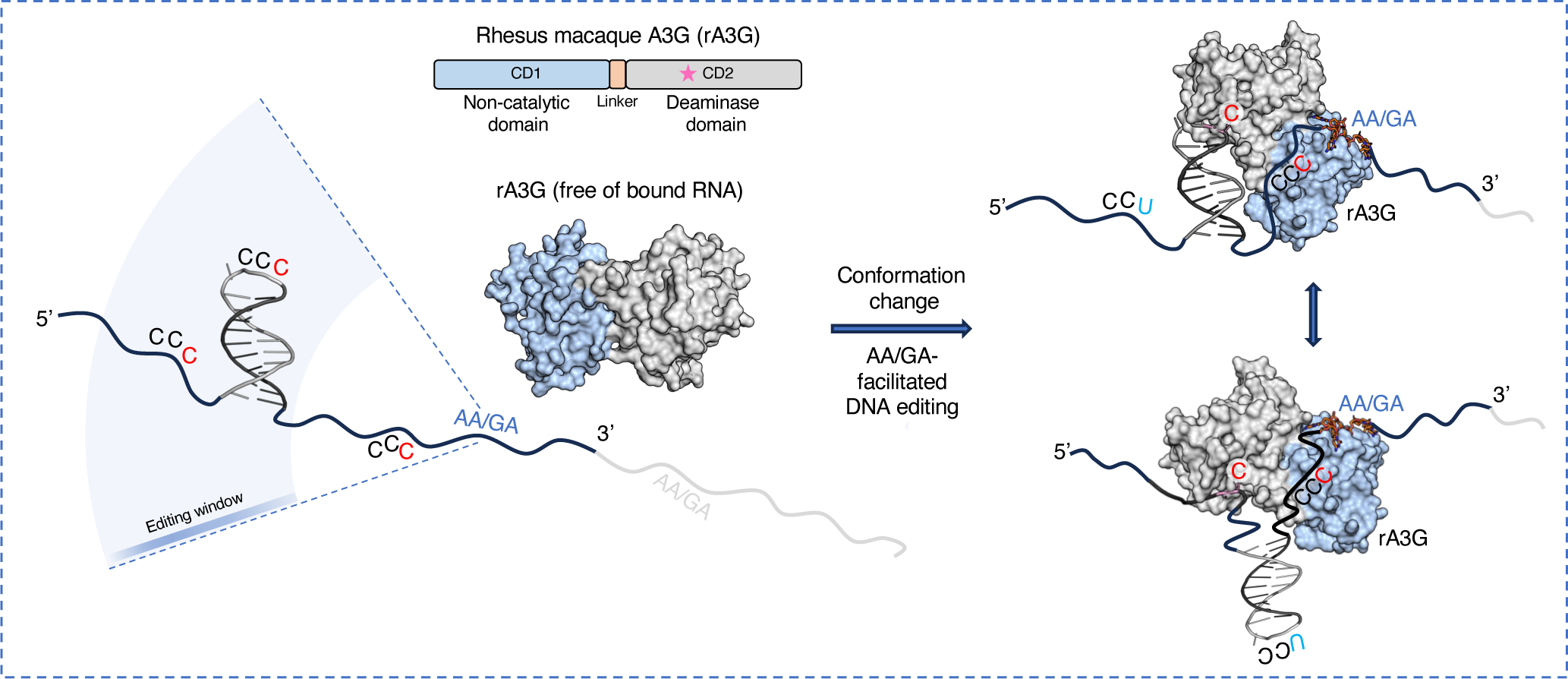
AA/GA-facilitated DNA editing by rA3G. rA3G binds the exposed AA/GA motifs on DNA and edits multiple CCCs that are located (1) 5’ upstream and (2) within its editing window (outside of the minimal effective distances required for linear or for hairpin DNA). The diagram depicts the fate of three editing sites: the target-C on the right remains unedited, whereas both middle and left target-Cs are edited. The target-C on the right could also be edited if rA3G binds to the AA/GA motif in light gray color.

rA3G also carries out AA/GA independent DNA editing at a reduced efficiency, which requires further study. The editing on ssDNA devoid of AA/GA may be carried out by the action of CD2 without much contribution from CD1, or there could be other DNA binding sites present on CD1 in either sequence specific or non-specific manner. Indeed, the editing level of ssDNA devoid of AA/GA is similar to those by individual CD2 protein alone (Prochnow2007Nature). However, the editing efficiency of AA/GA-independent editing significantly increases as the DNA length 3’-downstream of the CC<u>C</u> motif increases (**Fig. 1h**, **Fig. 3i**), suggesting that the increase of the length of the ssDNA can enhance DNA binding by CD1 to facilitate the substrate capturing and target-C deamination. Additionally, rA3G mutants disrupting the AA/GA specific binding are shown to retain AA-independent DNA editing (**Fig. 3g, 3i**).

We consider the following two factors as important for AA/GA-facilitated DNA editing: (1) differential binding affinity to purine and pyrimidine motifs on DNA and (2) a physical barrier between the catalytic cavity and the motif binding pocket. Variations in both binding affinity and flexibility degree between the two domains could influence the outcome of enzyme activity among other A3G homologs. Additionally, multiple factors could potentially shape the contribution of AA/GA-facilitated editing under physiologically relevant situations. Due to the largely shared binding interactions between AA/GA-containing ssDNA and ssRNA, mutants defective in binding to AA/GA ssDNA are also defective in binding to AA/GA ssRNA^33^. Cellular and viral ssDNA binding proteins could compete with A3G in binding to ssDNA^63–66^. Furthermore, temporary formed DNA secondary structures may cause deviation from expected productive configuration for AA-facilitated editing^7,67,68^. Finally, selection pressure in living cells may promote certain mutations over others and shape the eventual mutational outcome. Further investigation is needed to determine the mechanisms of CD1/CD2 cooperativity, and the outcome of target-C editing/mutation carried out by A3G in vivo. Structural study of other double-domain APOBECs can also offer valuable molecular insights into the cooperative mechanisms that have evolved during conflicts between host restriction factors and retroviruses.

## Methods

### Protein expression and purification

A soluble variant of rhesus macaque APOBEC3G (rA3G, accession code: AGE34493) with a replacement in the N-terminal domain loop 8 (^139^CQKRDGPH^146^ to ^139^AEAG^142^, designated as rA3G^R8^) was constructed in the pET-sumo vector (Thermo Fisher Scientific). This construct generated a SUMO fusion with an N-terminal 6xHis tag and a PreScission protease cleavage site. Protein expression and purification followed previously published protocols^33^. In brief, *E. coli* Rosetta™(DE3) pLysS cells expressing rA3G were cultured at 37°C in LB medium with 50 μg/ml kanamycin. Temperature was lowered to 16°C when OD_600_ reached ∼0.3.

Protein expression was induced by 0.1 mM IPTG when OD_600_ reached 0.7 to 0.9. After overnight growth at 16°C, cells were harvested by centrifugation. The resulting cell pellet was lysed in buffer A (25 mM HEPES at pH 7.5, 500 mM NaCl, 20 mM MgCl_2_, 0.5 mM TCEP, and 60 μg/ml RNase A) using sonication. The protein purification process included Ni-NTA agarose chromatography, RNase A/T1 treatment and PreScission protease cleavage, heparin chromatography, and size-exclusion chromatography. The final protein samples were quantified, verified for purity, and stored in buffer B (50 mM HEPES at pH 7.5, 250 mM NaCl, and 0.5 mM TCEP) at −80°C until use. Sequences of all mutant constructs were verified by Sanger sequencing (Azenta Life Sciences). Mutant proteins were purified using the same protocol.

### Electrophoretic mobility shift assay (EMSA)

DNA labeled with 5’ 6-FAM at 10 nM was titrated by rA3G in 20 μl reaction volume containing 50 mM HEPES pH 7.5, 250 mM NaCl, 1 mM DTT, 0.1 mg/ml recombinant albumin (New England Biolabs), 0.1 mg/ml RNase A (QIAGEN), and 10% glycerol. Reaction mixtures were incubated on ice for 10 min and analyzed by 8% native PAGE in 4 °C. A solution with acrylamide:bis-acrylamide ratio of 72.5:1 was used in preparing 8% native gels. AmershamTM TyphoonTM Biomolecular Imager (GE Healthcare) was used to visualize gel images. ImageQuant TL (GE Healthcare) was used for image quantification. Dissociation constant K_D_ was calculated using GraphPad Prism version 8.0.0 for Windows. Three independent experiments were carried out for each DNA molecule.

### In vitro UDG-dependent deaminase activity assay

DNA and RNA oligonucleotides were synthesized by Integrated DNA Technologies. Hairpin DNA substrates are annealed overnight in the DNA annealing buffer (10 mM Tris at pH 8, 50 mM NaCl), and their size exclusion chromatography (SEC) elution profiles were checked to ensure no self-dimer formation.

DNA deamination activity assays were performed as described^31^ with minor modifications. Reactions (20 μl) containing the purified protein (rA3G_R8_ or indicated mutants, with specified concentrations) and individual DNA substrates (with indicated sequences and concentrations) were incubated at 37°C for an indicated duration in DNA deamination buffer [25 mM HEPES at pH 7, 250 mM NaCl, 1 mM DTT, 0.1% Triton X-100, 0.1 mg/ml recombinant albumin (New England Biolabs), and 0.1 mg/ml RNase A (QIAGEN)]. Reactions were stopped by heating to 90 °C for 5 minutes. Uracil was removed by uracil DNA glycosylase (0.025 U/μl, New England Biolabs) at 37 °C for 15 minutes, followed by abasic site hydrolysis at 90°C for 10 minutes in 0.1 M NaOH. Reactions were mixed with equal volume of 2X gel loading buffer (95% formamide, 25 mM EDTA) and heated to 95 °C for 5 minutes. DNA fragments was separated on 20% denaturing acrylamide gel (5% crosslinker, 7 M urea, 1X TBE buffer) using Criterion™ cell apparatus (Bio-Rad) at 300 V for 40 to 60 minutes. For unlabeled DNA, gels were stained with 1X SYBR™ Gold Nucleic Acid Gel Stain (Thermo Fisher Scientific) for 10 minutes. Gel images were visualized by Typhoon^TM^ Biomolecular Imager (Cytiva) and quantified by ImageQuant TL image analysis software (Cytiva). The percent product formation was calculated by dividing the intensity of the lower product band by the sum of the intensities of the product and substrate bands. RNA competition experiments were conducted with both RNA and DNA present in the reaction mixture, following the same protocol.

### Crystal growth, data collection, Structure determination, and analysis

rA3G^R8^ carrying the inactive mutation E259A (rA3G^R8/E259A^) was purified using the same protocol as described above. The rA3G^R8/E259A^-DNA complexes were prepared by mixing protein (4 mg/ml) with DNA at 1 to 1 molar ratio. After incubating on ice for 1 hour, precipitation was removed by centrifugation (21,000×g, 2 minutes, 4°C). Initial screening was conducted using the sitting-drop vapor diffusion method with the ARI Crystal Gryphon Robot (ARI) and crystallization screening kits (QIAGEN) at 18°C. Crystallization hits were further optimized using the hanging-drop vapor diffusion method at 18°C. High-quality crystals of the rA3G^R8/E259A^-AA DNA complex were obtained with a reservoir solution consisting of 0.1 M Bis-Tris Propane at pH 7.3, 0.2 M Na/KPO_4_, and 18% PEG 3350. Similarly, high-quality crystals of the rA3G^R^^8/E259A^-GA DNA complex were obtained with a reservoir solution consisting of 0.1 M Bis-Tris Propane at pH 7.5, 0.2 M Na/KPO_4_, and 16% PEG 3350. These crystals were transferred to synthetic mother liquor supplemented with suitable amounts of glycerol for cryoprotection and then flash-cooled in liquid nitrogen. X-ray diffraction data were collected at the Advanced Photon Source (GM/CA@APS, Argonne National Laboratory) beamline 23ID-D and at the Advanced Light Source (ALS, Lawrence Berkeley Laboratory) beamline 5.0.3.

Data were processed by automated data processing pipelines at the beamlines. Initial phase information was obtained by molecular replacement method with PHENIX using the rA3G crystal structures (PDB 7UU4 or 8EDJ) without bound RNA. DNA was built manually in COOT. The structural models were refined using PHENIX and modified with COOT. Data collection statistics and refinement parameters are summarized in Table 1. Hydrogen bonding predictions were done by QtPISA. Structure images were prepared with PyMOL.

**Table 1.**
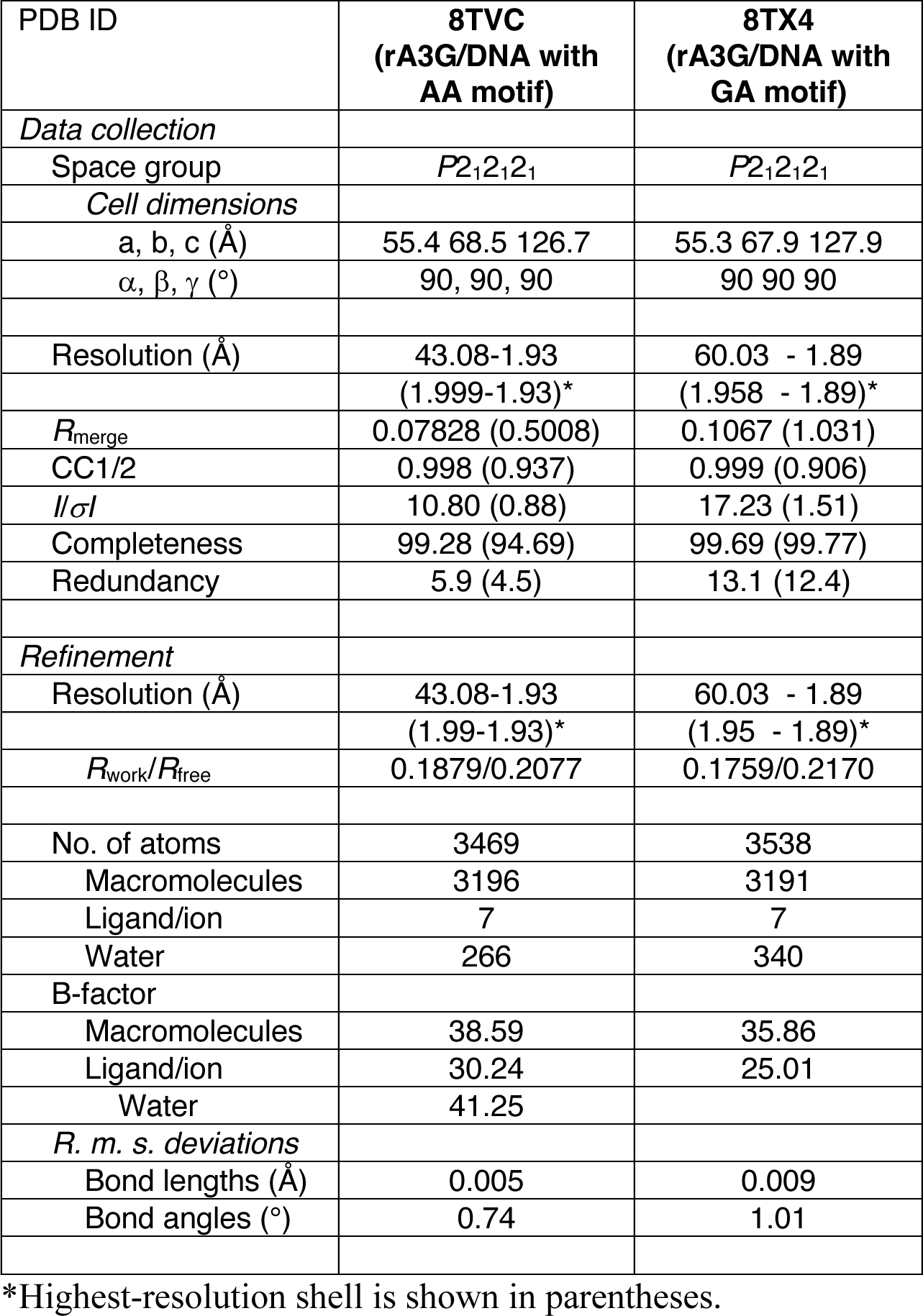
Crystallographic data collection and refinement statistics.

## Data availability

Atomic coordinates and structure factors have been deposited in the PDB database under accession codes 8TVC (rA3G^R^^8/E259A^ in complex with DNA 5’-CAATC) and 8TX4 (rA3G^R^^8/E259A^ in complex with DNA 5’-TGAT).

## Acknowledgement

We thank Phuong Pham and Malgorzata Jaszczur for advice on functional biochemical analyses.

Beamlines of GM/CA@APS have been funded by the National Cancer Institute (ACB-12002) and the National Institute of General Medical Sciences (AGM-12006, P30GM138396). The ALS-ENABLE beamlines are supported in part by the National Institutes of Health, National Institute of General Medical Sciences, grant P30 GM124169. This work is supported by the NIH grant R01 AI150524 to X.S.C.

## Author contributions

H.Y. and X.S.C. designed the experiments. H.Y. performed crystallization, data collection, structural determination, and functional biochemical analyses. J.P. performed structural work and refinements. H.Y. and X.S.C. wrote the manuscript. All authors commented on the manuscript.

