## Supplemental Figures for "Molecular mechanism for regulating APOBEC3G DNA editing function by the non-catalytic domain"

[1]Molecular and Computational Biology, Departments of Biological Sciences, University of Southern California, Los Angeles, CA 90089, USA. [2]Texas Biomedical Research Institute, San Antonio, TX 78227, USA, [3]Department of Microbiology, Immunology and Molecular Genetics, and [4]California NanoSystems Institute, University of California, Los Angeles, CA90095, USA. [5]Genetic, Molecular and Cellular Biology Program, Keck School of Medicine, [6]Norris Comprehensive Cancer Center, [7]Center of Excellence in NanoBiophysics, University of Southern California, Los Angeles, CA 90089, USA.

*To whom correspondence should be addressed.

**This PDF file contains:**

- Supplementary Figure 1 - 9

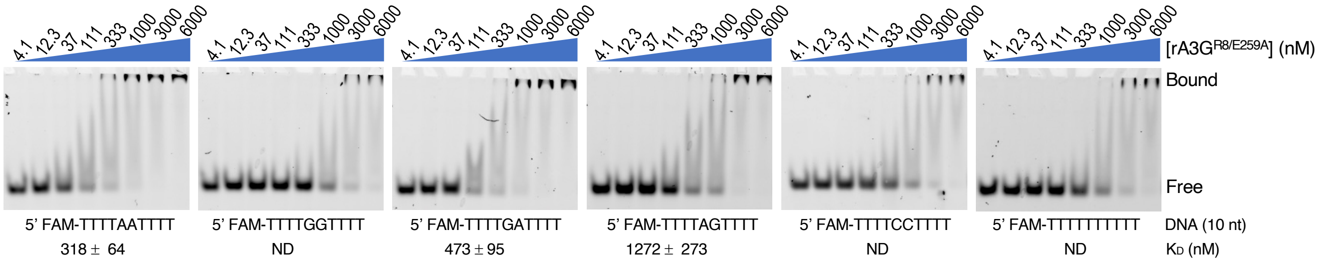


**Supplementary Figure 1**

Binding of rA3G$^{R8/E259A}$ to synthesized DNA molecules was visualized through electrophoretic mobility shift assay (EMSA). DNA (5' FAM-TTTTNNTTTT, NN = AA, GG, GA, AG, CC, or TT, 10 nt) at a fixed concentration of 10 nM was incubated with rA3G$^{R8/E259A}$ at various concentrations (4.1, 12.3, 37, 111, 333, 1000, 3000, or 6000 nM). The estimated $K_D$ values as mean values ± SD are listed below individual representative EMSA gel images. n = 3 independent experiments.

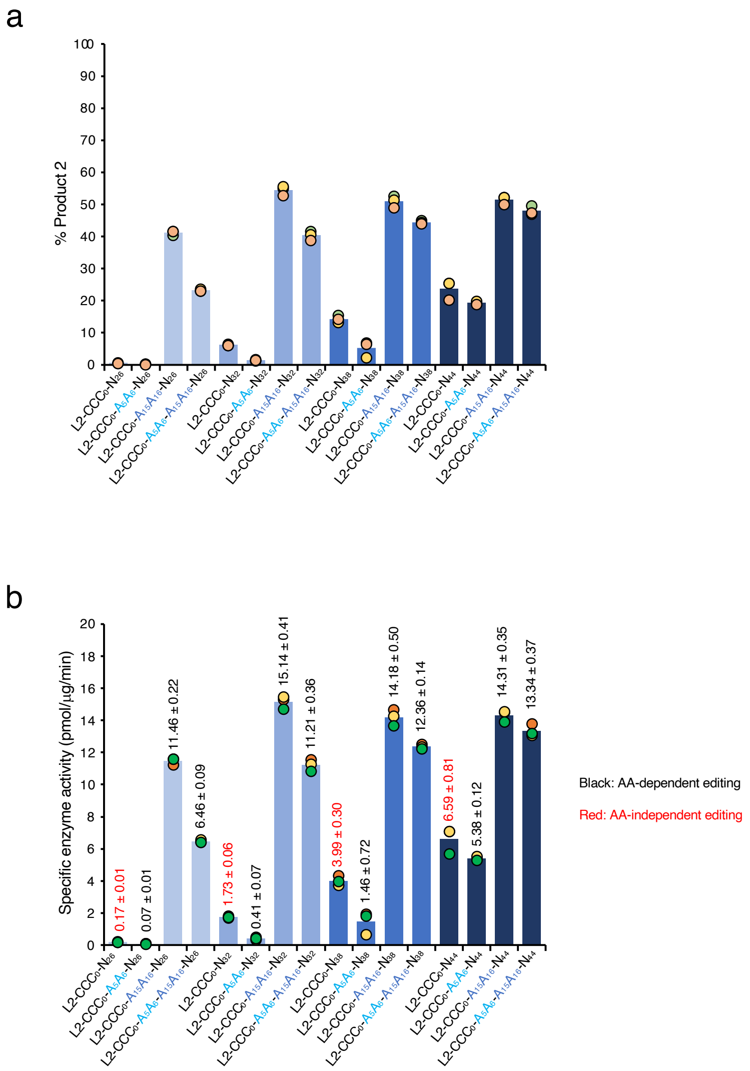


**Supplementary Figure 2**

Plots depicting the %product 2 formation (**a**) and specific enzyme activity (**b**) were derived from the data in **Fig. 1h**. Deamination reactions were carried out with 50 nM rA3G$^{R8}$ and 500 nM substrates in 20 µl reaction volume for 8 min at 37 °C, and gels were stained with SYBR Gold. The following formular was used to calculate specific enzyme activity, assuming that the product amount was 0 at time point 0. Specific enzyme activity = %Product*0.01*500000 (pM)*0.00002 (L)/0.045 (µg)/8 (min). The unit of the specific enzyme activity is pmol/µg/min.

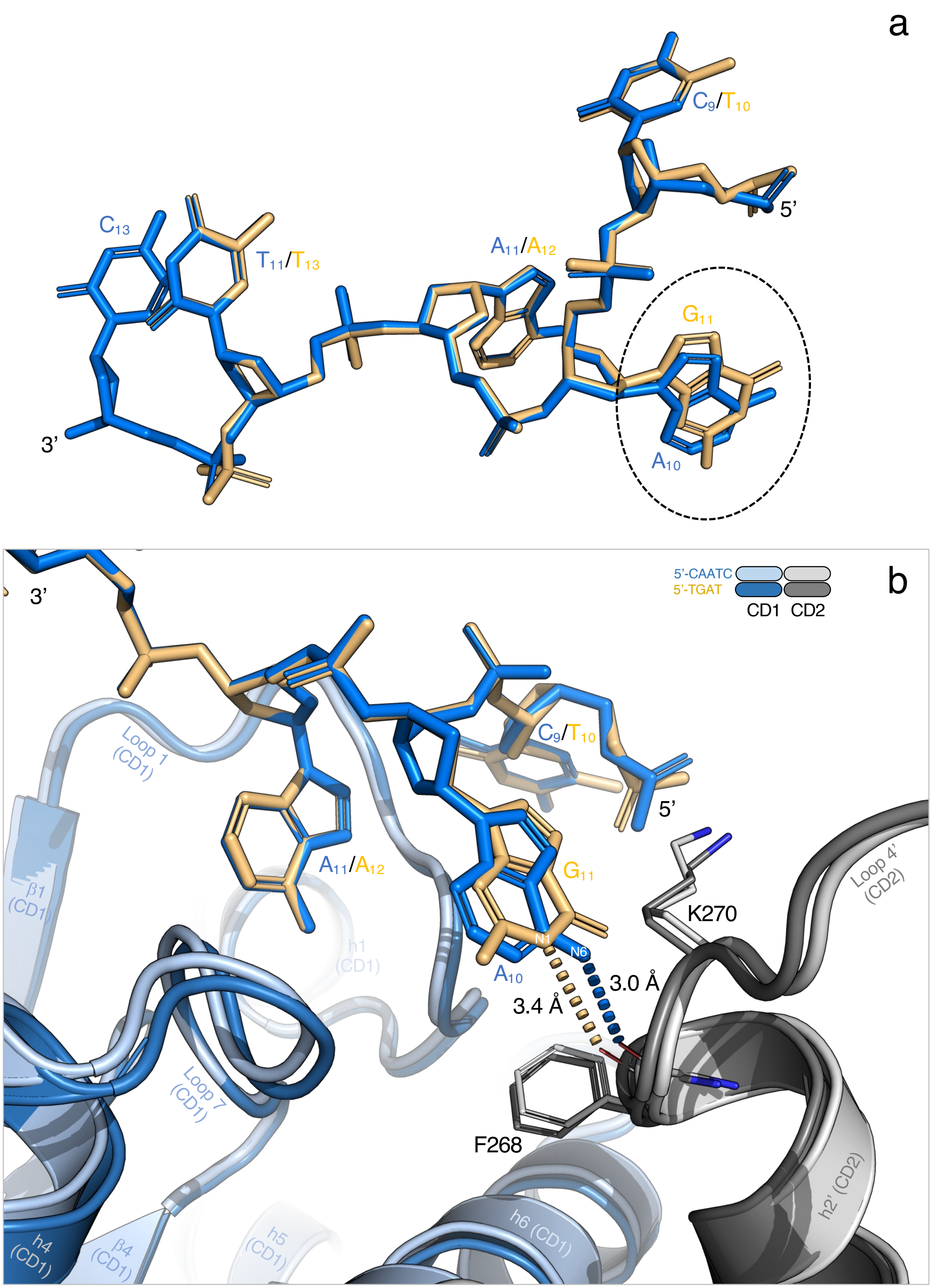


**Supplementary Figure 3**

(**a**) Superimposition of the two DNA molecules is depicted in stick representation, 5’-$C_9A_{10}A_{11}T_{12}C_{13}$ (in blue stick) and 5’-$T_{10}G_{11}A_{12}T_{13}$ (in yellow stick), revealing well-aligned bases with only a slight difference between the positions of $A_{10}$ and $G_{11}$ (represented by blue and yellow sticks, respectively, inside a black dotted circle). (**b**) Superimposition of the two DNA-rA3G$^{R8/E259A}$ complexes, showing the hydrogen bond between N6 of $A_{10}$ and the main-chain carbonyl group of F268 (indicated by a blue dashed line, 3.0 Å), and the hydrogen bond between N1 of $G_{11}$ and the main-chain carbonyl group of F268 (indicated by a yellow dashed line, 3.4 Å).

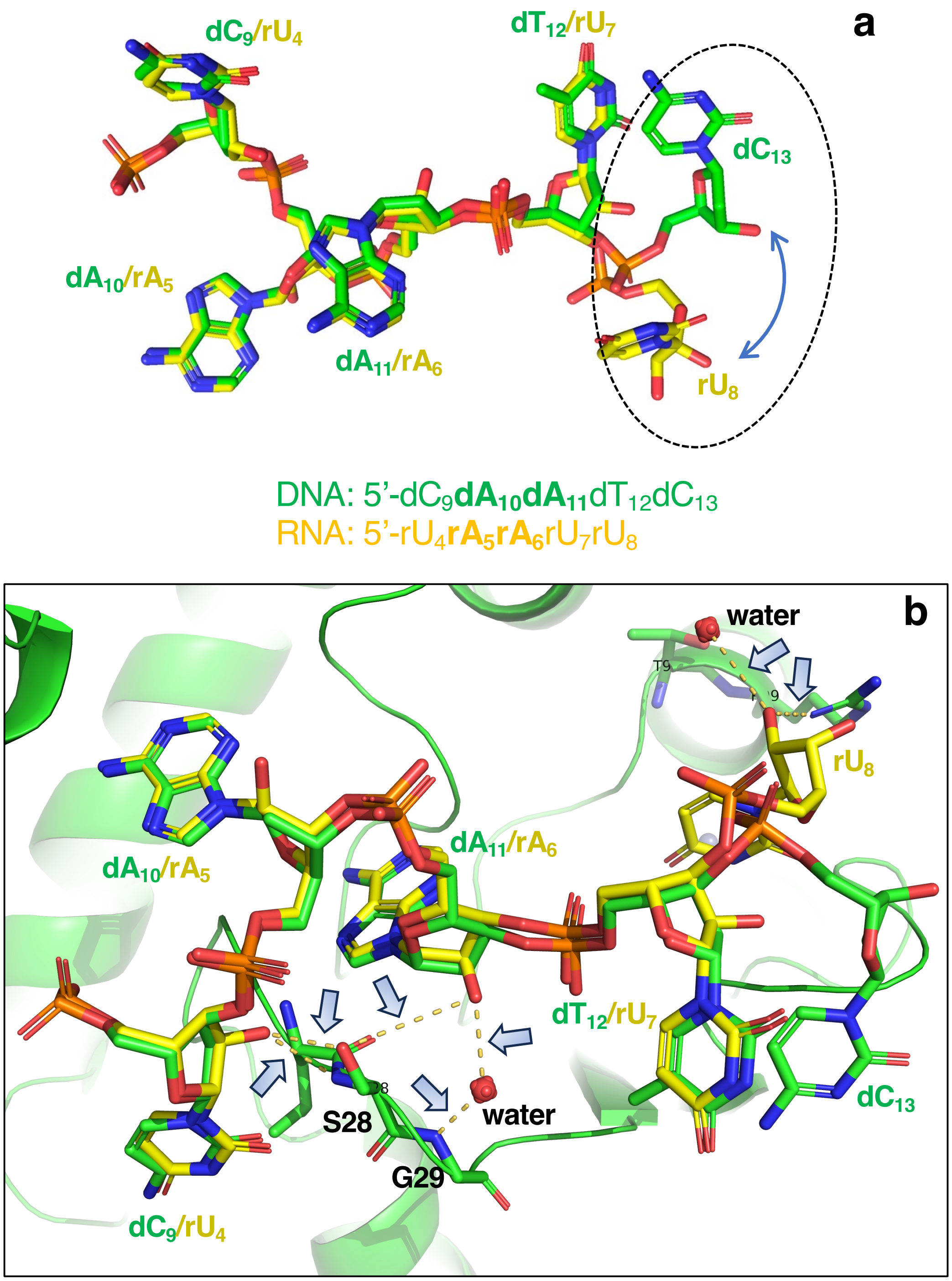


**Supplementary Figure 4**

(**a**) Superimposition of the DNA and RNA molecules is depicted in stick representation, 5'-$dC_9dA_{10}dA_{11}dT_{12}dC_{13}$ (in green stick) and 5'-$rU_4rA_5rA_6rU_7rU_8$ (in yellow stick, PDB ID 7UU4[33]), revealing well-aligned bases with only the difference between the positions of $dC_{13}$ and $rU_8$ (represented by green and yellow sticks, respectively, inside a black dotted circle). (**b**) Superimposition of the DNA-rA3G$^{R8/E259A}$ and RNA-rA3G$^{R8/E259A}$ complexes, showing multiple RNA-specific hydrogen bonds (indicated by yellow dashed lines and blue arrows).

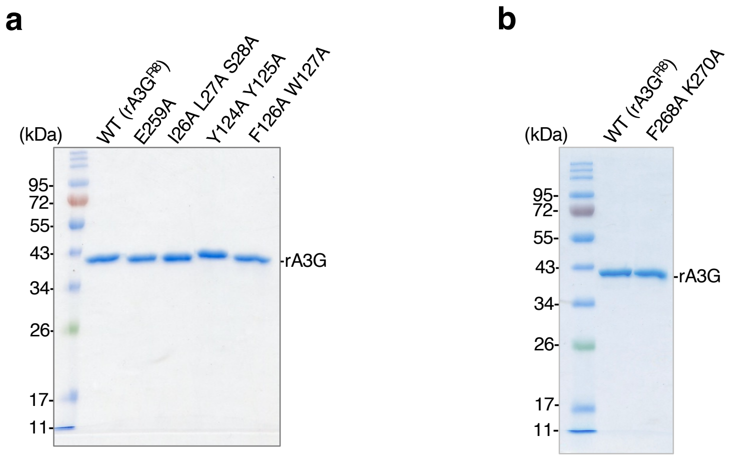


**Supplementary Figure 5**

SDS-PAGE gel images showing the purified rA3G$^{R8}$ (wild-type equivalent) and the mutants used in **Fig. 3** and **Fig. 4**. Each lane was loaded with a 2 μl protein sample at a concentration of 10 μM (~900 ng protein).

**a** Hairpin DNA 1 set (stem length: 10 bp)

| | |
|---|---|
| Substrate length | 45 nt |
| Stem length | 10 bp |
| $T_m$ | 75.4 to 76.9 °C |
| Running buffer | 25 mM HEPES, pH 7.5, 250 mM NaCl |

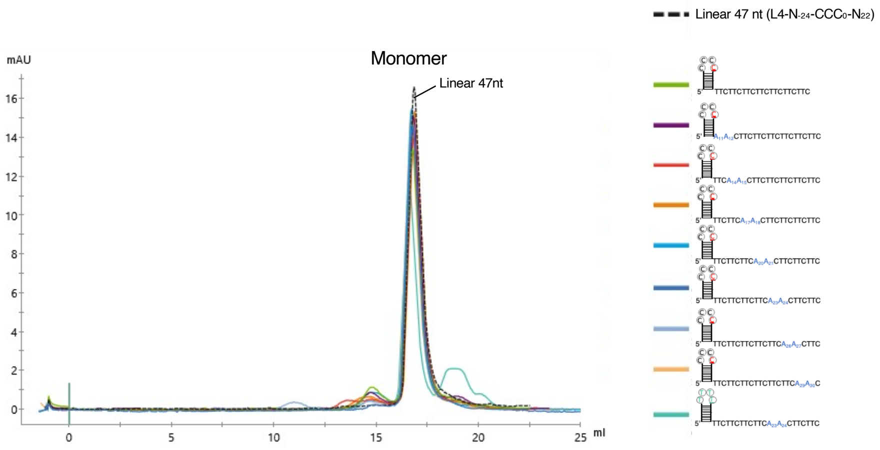


**b** Hairpin DNA 2 set (stem length: 4 bp to 12 bp)

| | |
|---|---|
| Substrate length | 33 nt to 49 nt |
| Stem length | 4 bp to 12 bp |
| $T_m$ | 61.8 to 77.4 °C |
| Running buffer | 25 mM HEPES, pH 7.5, 250 mM NaCl |

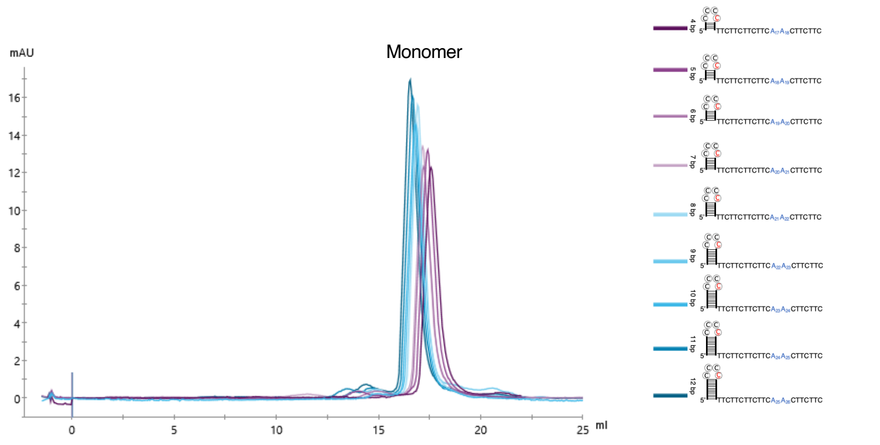


**Supplementary Figure 6**

Size exclusion chromatography (SEC) elution profiles of the annealed hairpin DNA substrates, showing the annealed hairpin DNA molecules are monomeric under experimental conditions.

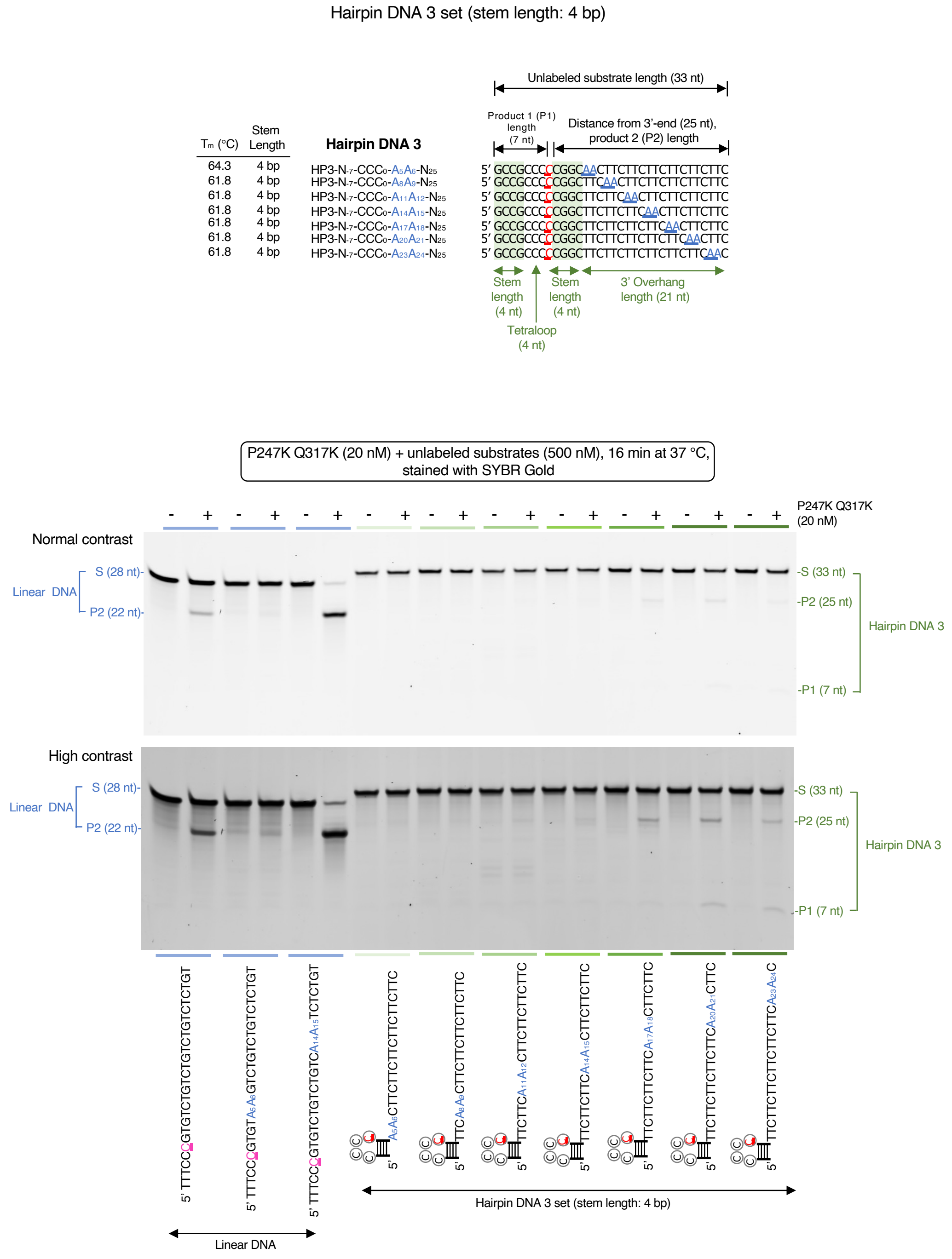


### Supplementary Figure 7

Deamination activity was monitored on hairpin DNA 3 substrates carrying a 4-bp hairpin stem, a tetraloop CCCC, and a 3'overhang. A single AA motif was placed at various locations on the 3' overhang. Three linear DNA substrates were included as linear DNA controls. The results show that hairpin DNA with the stem length of 4 bp are poorly edited under the experimental conditions.

Hairpin DNA 4 set (stem length: 11 bp)

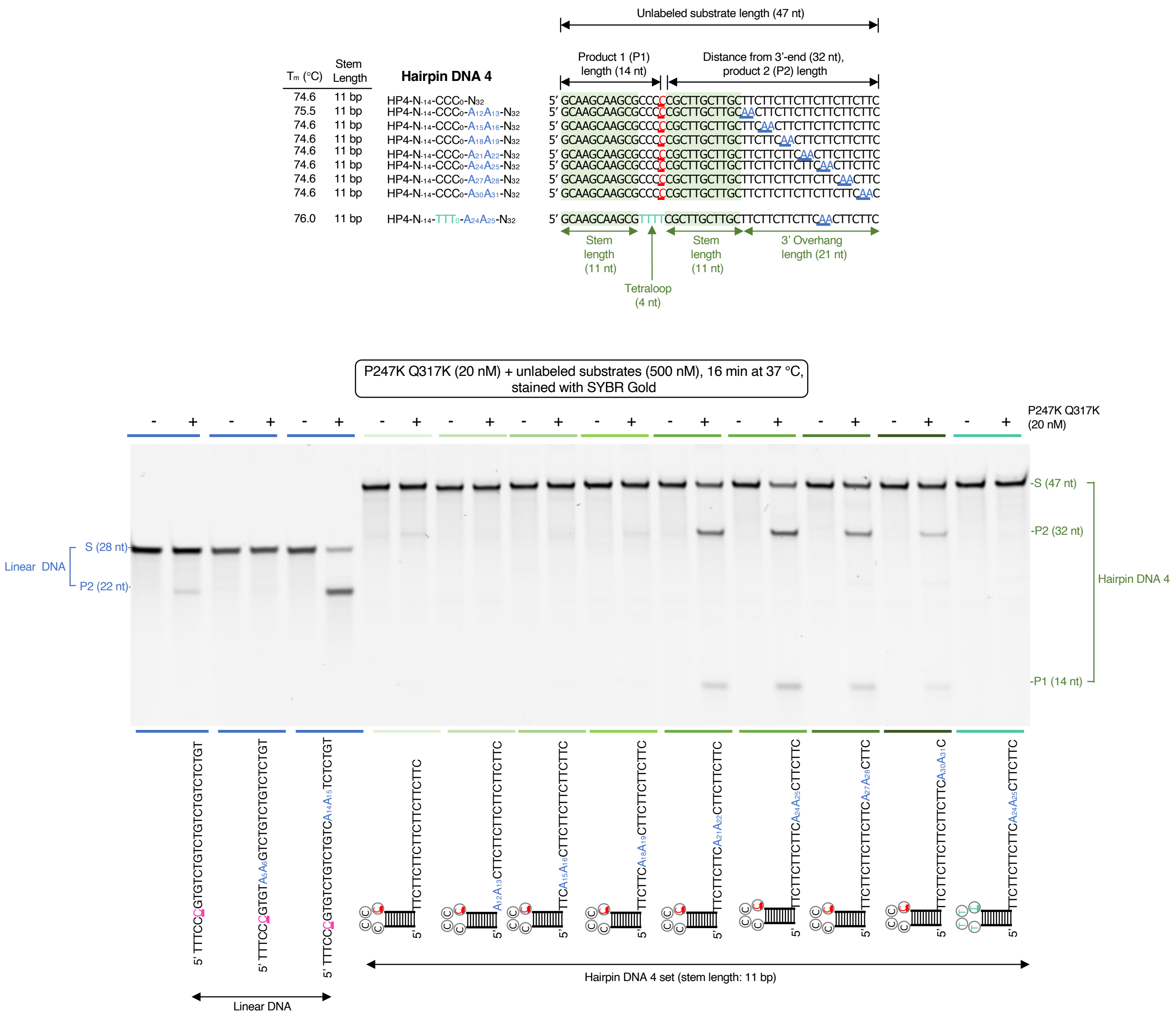


### Supplementary Figure 8

Deamination activity was monitored on hairpin DNA 4 substrates carrying a 11-bp hairpin stem, a tetraloop CCCC, and a 3'overhang. A single AA motif was placed at various locations on the 3' overhang. A negative control with 5'-TTTT in the hairpin loop and a single $A_{24}A_{25}$ motif in the 3'overhang was included. Three linear DNA substrates were also included as linear DNA controls. The results show that the top three edited hairpin DNA with the stem length of 11 bp are those carrying a single AA motif $A_{21}A_{22}$, $A_{24}A_{25}$, or $A_{27}A_{28}$.

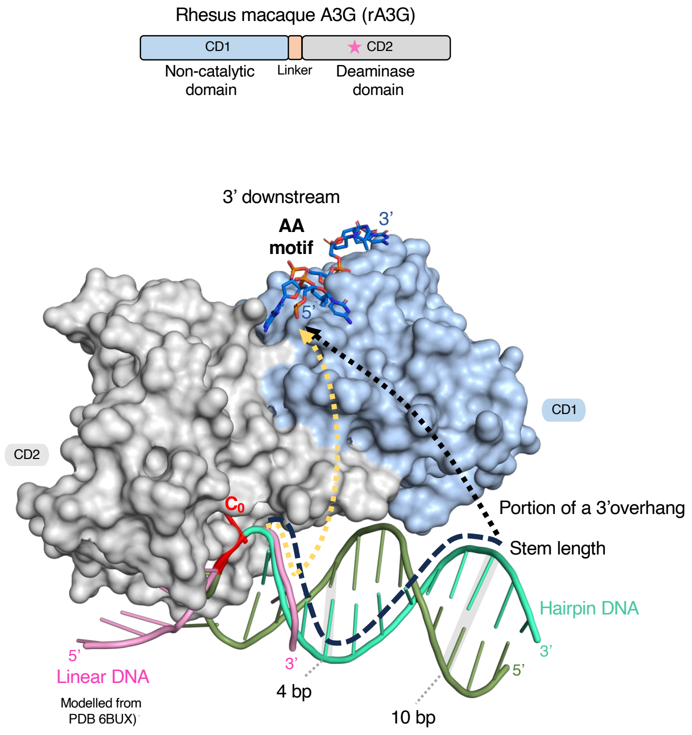


- The yellow path is for linear DNA
- The black path is for hairpin DNA

**Supplementary Figure 9**

This model illustrates the potential connection paths between the CCC motif bound to CD2 and the AA motif bound to CD1 (3' downstream) in both linear DNA (indicated by a yellow dotted line) and hairpin DNA (indicated by a black dotted line). When CCC motif is on the loop of a stem-loop structure, the distance from CCC connecting to AA (3' downstream) is the sum of the stem length and the portion of a 3' overhang. Consequently, the principle governing the spatial requirement is consistent across both linear ssDNA and stem-loop DNA, despite differences in the number of nucleotides between the target cytosine $C_0$ and AA motif.
